# Increasing the representation of minoritized youth for inclusive and reproducible brain-behavior associations

**DOI:** 10.1101/2024.06.22.600221

**Authors:** Jivesh Ramduny, Lucina Q. Uddin, Tamara Vanderwal, Eric Feczko, Damien A. Fair, Clare Kelly, Arielle Baskin-Sommers

## Abstract

Population neuroscience datasets allow researchers to estimate reliable effect sizes for brain-behavior associations because of their large sample sizes. However, these datasets undergo strict quality control to mitigate sources of noise, such as head motion. This practice often excludes a disproportionate number of minoritized individuals. We employ motion-ordering and motion-ordering+resampling (bagging) to test if these methods preserve functional MRI (fMRI) data in the Adolescent Brain Cognitive Development Study (*N*=5,733). Black and Hispanic youth exhibited excess head motion relative to data collected from White youth, and were discarded disproportionately when using conventional approaches. Both methods retained more than 99% of Black and Hispanic youth. They produced reproducible brain-behavior associations across low-/high-motion racial/ethnic groups based on motion-limited fMRI data. The motion-ordering and bagging methods are two feasible approaches that can enhance sample representation for testing brain-behavior associations and fulfill the promise of consortia datasets to produce generalizable effect sizes across diverse populations.

## Main

Population neuroscience datasets made available through consortia efforts provide unprecedented opportunities for researchers to relate individual differences in brain functions (e.g., functional connectivity) to individual differences in behavior (e.g., general psychopathology, cognitive ability) with sufficient statistical power^1–3^. These datasets have accelerated scientific discovery, improved reproducible research practices, and democratized the field by reducing the resources required to do research^4–7^. Given the increasing cost of acquiring an MRI scan (US$500-US$1,000/HR), combined with the fact that many researchers do not have access to an MRI scanner, particularly in the Global South, these consortia datasets (*N* ≳ 1,000) purportedly have allowed researchers to estimate reliable effect sizes of brain-behavior associations because of their large sample sizes^1^. However, consortia datasets do not necessarily capture diverse populations, limiting our ability to yield brain-behavior relationships that are generalizable across diverse sociodemographic characteristics^8,9^.

Based on the 2021 US Census^10^, 59.3% of the population is White and not Hispanic or Latino, 18.9% is Hispanic, and 13.6% is Black or African American. However, according to the 2018-2021 National Institutes of Health (NIH) Research Condition and Diseases Categorization (RCDC) Inclusion Statistics Report^11^, White participants are overrepresented in human neuroscience studies with a median proportion of ∼70% across all NIH funded neuroscience studies. The lack of participant inclusivity is evident in many consortia datasets such as the Human Connectome Project [HCP] (*N*≈1,200) and UKBiobank [UKB] (*N*≈60,000). The HCP and UKB respectively include ∼76% and ∼95% White participants^8^. However, the Adolescent Brain Cognitive Development^SM^ Study (ABCD Study^®^; *N*≈12,000) followed the American Community Survey and National Center for Education Statistics to represent ∼52% White, ∼15% Black, and ∼20% Hispanic youth^12^. The ABCD Study aimed to recruit youth longitudinally by sampling the sociodemographic variations in the US population^12^.

Not only are minoritized individuals underrecruited in many neuroscience studies, but they also tend to disproportionately be excluded following standard pre-processing protocols typically used for MRI data analysis. To minimize variations in MRI data acquisition across consortia sites, researchers tend to apply strict quality control strategies to mitigate potential biases. One quality control strategy is to address the potential impact of participant head motion^1^. Head motion can significantly inflate functional connectivity estimates across a diverse range of populations^13–19^. This issue is exacerbated in studies of youth due to the presence of heightened in-scanner motion relative to adult populations and a robust negative correlation between head motion and age^13,20–23^. Multiple strategies already exist to correct for motion^15,16,24–26^, but none succeed in removing the influence of motion completely. To reduce the impact of head motion in brain-behavior association studies, data exclusion practices are commonly applied that result in discarding large numbers of participants^1^. These practices inadvertently tend to exclude minoritized youth disproportionately, due to lower adherence to mainstream protocols and a lack of culturally informed strategies to engage and retain racial/ethnic minorities in neuroimaging research, particularly those from underprivileged backgrounds^8,27,28^.

Here, we built on previous work^29^ by applying two head motion mitigation methods — motion-ordering and motion-ordering+resampling (i.e., bagging) — for estimating functional connectivity to maximize participant inclusion from “high-motion” minoritized youth (see **Online Methods** for rationale). We aimed to retain as many minoritized youth as possible using their motion-limited fMRI data for establishing inclusive and reproducible brain-behavior associations from the ABCD Study^30^. We selected the ABCD Study for its census distribution of races and ethnicities including White (64%), Black (14%), and Hispanic (22%) youth with adequate sample sizes. To assess the validity of the two methods, whole-brain brain-behavior associations between functional connectivity and cognitive performance and between connectivity and psychopathology were estimated for a sample that included “high-motion” youth across the 3 racial/ethnic groups, who otherwise would have been discarded using traditional criteria for data exclusion based on excessive head motion.

## Results

Complete demographic, behavioral, and imaging data from 5,733 youth were obtained from the ABCD Study Baseline Release (9-10 years)^30^. Youth categorized as belonging to one of the 3 racial/ethnic groups, White, Black, and Hispanic were included in the present study; youth who belonged to other racial/ethnic categories were not included due to their limited sample sizes (**Online Methods**). For each youth, a whole-brain parcellation scheme was applied to extract their resting-state fMRI timeseries concatenated across a maximum of 4 runs. Functional connectivity was derived from the fMRI timeseries by computing and Fisher *z*-transforming the Pearson correlation coefficient (*r*) between all possible pairs of ROIs using an extended Gordon parcellation scheme^31,32^ to construct their 352 x 352 functional connectivity matrices. This pipeline is referred to as the “standard approach”.

The motion-ordered and bagging methods were then performed by applying a frame censoring procedure known as scrubbing^14,18,33,34^ to identify and remove motion-corrupted timepoints (*T*) in the fMRI timeseries (see **Fig.1** for a schematic). Each *T* was identified with a head motion threshold of mean framewise displacement (FD)^18^ > 0.20 mm. For each *T*, one preceding (*T* – 1) and two succeeding (*T* + 1, *T* + 2) timepoints were censored in the fMRI timeseries to minimize the presence of residual motion and micro head movements. For each participant, we created subsets of the scrubbed timepoints that matched a predefined threshold referred to as minTP to prevent participants from having varying number of motion-limited timepoints in their timeseries. The scrubbed timeseries were ranked by their lowest FD values and the top minTP-matched timepoints were selected. For the motion-ordered method, the functional connectivity matrix of each participant was computed using the minTP-matched timepoints^29^. For bagging, 500 bootstrapped samples of size TP were generated *with* replacement for each participant using the minTP-matched timepoints^29^. Iteratively, the bagged functional connectivity matrix of each participant was recomputed. A mean bagged functional connectivity matrix also was generated for each participant by averaging the 500 connectivity matrices.

**Fig. 1.**
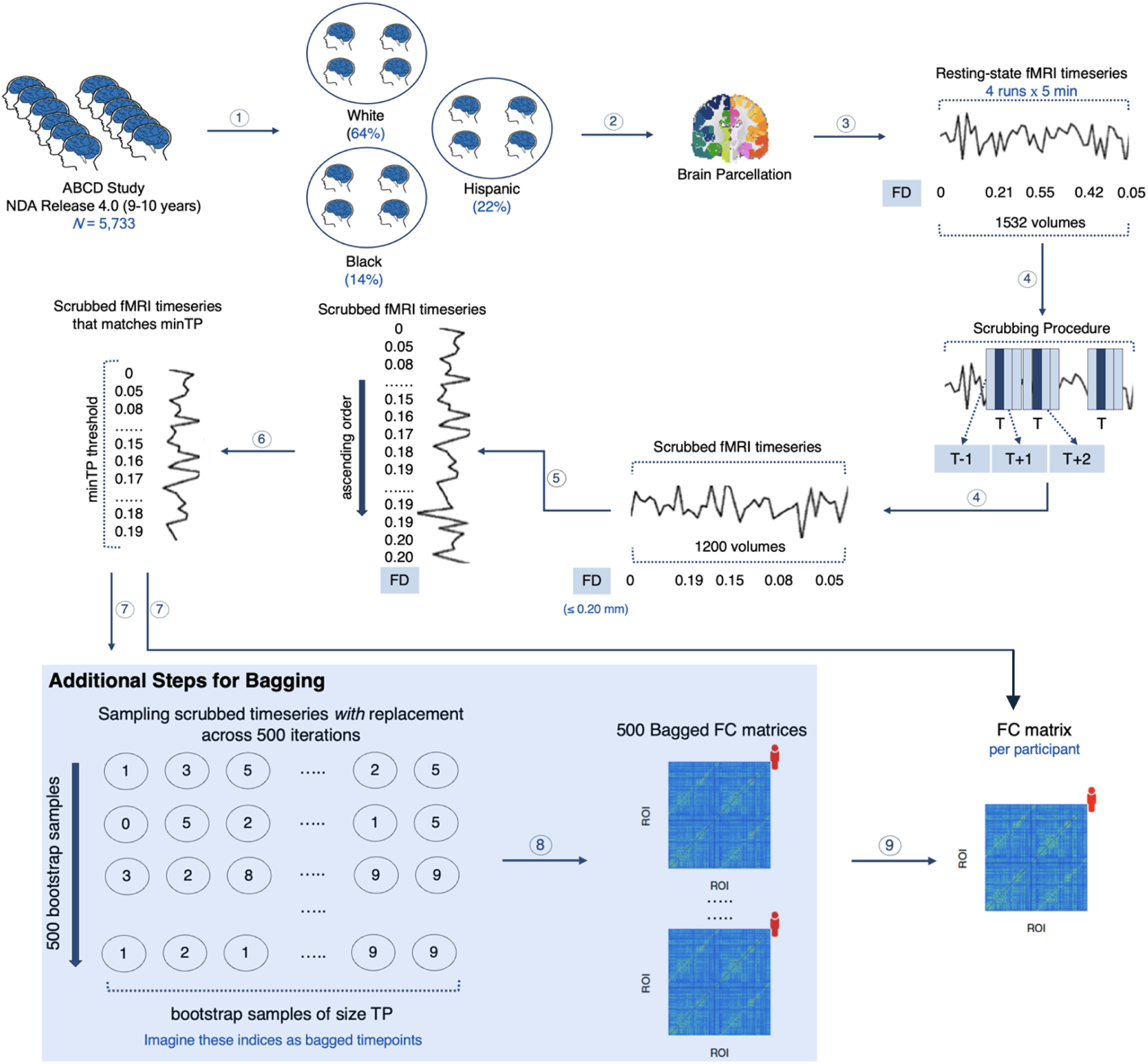
Motion-ordered and Bagging methods. 1. The participants in the ABCD Study were categorized into 3 racial/ethnic groups: White, Black, and Hispanic. 2. A whole-brain parcellation was applied to extract the resting-state fMRI timeseries of the youth concatenated across a maximum of 4 runs. 3. The framewise displacement (FD) was computed for every timepoint in the fMRI timeseries to quantify the amount of in-scanner motion for one volume relative to its preceding one. 4. Scrubbing was applied to identify and remove motion-corrupted timepoints (*T*) if their FD > 0.20 mm. For each *T*, one preceding (*T*) and two succeeding (*T* + 1, *T* + 2) timepoints were further censored. 5. For each youth, their scrubbed timeseries were ranked by their least FD values (0 → 0.20 mm). 6. A minTP threshold was imposed to prevent the youth from having different number of timepoints in their timeseries. 7. For the motion-ordered method, the functional connectivity (FC) of each participant was computed between all pairs of ROIs using Pearson correlation coefficient from the minTP-matched timepoints. For the bagging method, 500 bootstrapped timeseries samples of size TP were generated *with* replacement for each participant using the minTP-matched timepoints. 8. Iteratively, the functional connectivity (FC) between all pairs of ROIs was computed using Pearson correlation coefficient for the 500 bootstrapped samples. 9. A mean bagged functional connectivity (FC) matrix also was generated for each youth by averaging the 500 connectivity matrices. The motion-ordered and bagging methods are adapted from previous work^29^. The analysis code to this study can be found at https://github.com/JRam02/inclusivity.

Finally, the standard, motion-ordered, and bagged functional connectivity matrices for each participant were used to compute the associations between connectivity patterns and behavior for cognitive performance (NIH Toolbox) and externalizing and internalizing psychopathology (Child Behavior Checklist [CBCL]).

### Disproportionate exclusion of high-motion minoritized youth

MRI data exhibiting high levels of motion artifacts typically are discarded in brain-behavior association studies to minimize the impact of this nuisance variable, which can inflate effect size estimates. As expected, a strict head motion threshold, such as mean FD < 0.08 mm^1^, drastically reduced the sample sizes across racial/ethnic groups, relative to a liberal head motion threshold, such as mean FD > 0.50 mm (**Fig.2A**). However, there was a disproportionate decline in the sample sizes of the Black and Hispanic groups regardless of the mean FD threshold. This was due to a significant difference in mean FD across the 3 racial/ethnic groups, agreeing with previous evidence from the ABCD Study^35^. Minoritized youth exhibited greater head motion relative to White youth (**Fig.2B**): Kruskal-Wallis test, *H* (2) = 61.9, *P* < 0.001; White median meanFD = 0.16 mm, Black median meanFD = 0.20 mm, and Hispanic median meanFD = 0.19 mm. We observed a gradual disproportionate decrease in the temporal signal-to-noise ratio (tSNR) of the Black and Hispanic youth when liberal mean FD thresholds were applied (**Fig.2A**). There also was a significant difference in the number of scrubbed timepoints across the 3 racial/ethnic groups. Minoritized youth had fewer number of scrubbed timepoints in their motion-limited fMRI timeseries relative to White youth (**Fig.2A**): Kruskal-Wallis test, *H* (2) = 34.7, *P* < 0.001; White median scrubbed timepoints = 1,016, Black median scrubbed timepoints = 936, and Hispanic median scrubbed timepoints = 940. To retain as many low-/high-motion youth as possible and to maximize participant inclusivity across the 3 racial/ethnic groups, a minTP threshold of 100 motion-limited timepoints was applied.

**Fig. 2.**
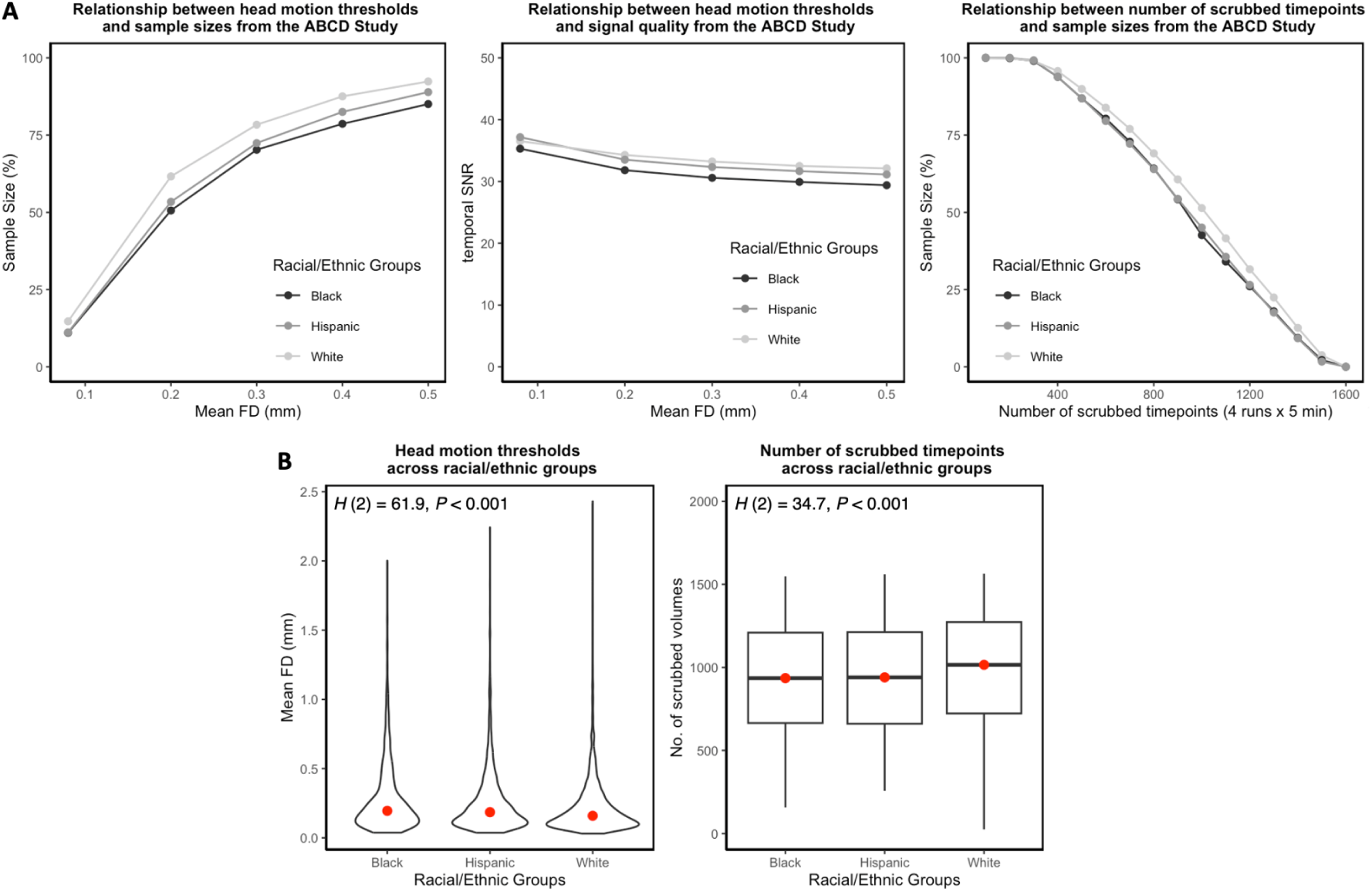
Impact of head motion across racial and ethnic groups in the ABCD Study. A. Left: Relationship between head motion thresholds and sample sizes. Head motion was quantified by mean framewise displacement (FD) such that mean FD ∈ {0.08, 0.20, 0.30, 0.40, 0.50}. Middle: Relationship between head motion thresholds and signal quality indexed by the temporal signal-to-noise ratio (tSNR). Mean FD ∈ {0.08, 0.20, 0.30, 0.40, 0.50}. Right: Relationship between number of scrubbed timepoints and sample sizes. Number of scrubbed timepoints ∈ {100, 200, 300, 400, 500, 600, 700, 800, 900, 1,000, 1,100, 1,200, 1,300, 1,400, 1,500}. B. Left: There was a significant difference in head motion across the 3 racial/ethnic groups. Right: There was a significant difference in the number of scrubbed timepoints across the 3 racial/ethnic groups.

### Standard brain-behavior associations from low-motion racial and ethnic groups

To compute the brain-behavior associations across the 3 racial/ethnic groups, partial Spearman’s Rank correlations (*R*_s_) were computed between functional connectivity and NIH Toolbox (measuring cognitive performance) as well as externalizing and internalizing CBCL scores, while treating sex assigned at birth and mean FD as covariates (**Online Methods**). We employed the standard method, which uses the full timeseries concatenated across a maximum of 4 resting-state runs from the low-motion youth. The low-motion youth were identified using a mean FD < 0.20 mm which retained a total of 3,342 participants, corresponding to ∼58% of the total sample size: White *N* = 2,266, Black *N* = 410, and Hispanic *N* = 666. There were significant differences in the behavioral distributions between the low-motion and high-motion youth across the 3 racial/ethnic groups (**Supplementary Figure 1**). The aim of this study was not to *compare* the brain-behavior relationships across the 3 racial/ethnic groups, but to generate associations that are inclusive and reproducible within races and ethnicities across the sample. The brain-behavior relationships were obtained from the strongest edge (*R*_s_) in the functional connectome after applying the Benjamini-Hochberg False Discovery Rate (q = 0.05)^36^ to correct for multiple comparisons across 61,776 edges (**Supplementary Figure 2**). While 17,854 edges shared a significant association with the NIH Toolbox, 65 edges yielded similar relationships with CBCL-externalizing. There was no edge that produced a significant association with CBCL-internalizing after correcting for multiple comparisons. Therefore, the brain-behavior analyses focused only on the NIH Toolbox and CBCL-externalizing measures.

We examined the reproducibility of the standard brain-behavior relationships as a function of *N* ranging from typical (*N* = 25) to large (maximum *N* of each group)^1^. For each low-motion racial/ethnic group and behavior, we computed a total of 11 intervals x 500 bootstrap *N* samples = 5,500 correlations. Random variation in a brain-behavior association across subsamples of the population can generate stronger associations with typical sample sizes by chance (i.e., sampling variability). Consequently, we would expect sampling variability to decrease and observe less inflated associations at larger sample sizes at a rate of √*N*^1^. We observed tightening of the 95% confidence interval (CI) as the sample sizes of the racial/ethnic groups grew from typical to large for both NIH Toolbox (**Table 1**) and CBCL-externalizing (**Table 2**). For NIH Toolbox, White 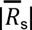 = 0.10, Black 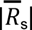 = 0.12, and Hispanic 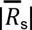 = 0.053. For CBCL-externalizing, White 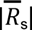 = 0.097, Black 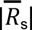 = 0.057, and Hispanic 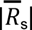 = 0.080.

**Table 1.** Correlations between functional connectivity and NIH Toolbox obtained from the standard, motion-ordered, and bagging methods as a function of sample size across the 3 low-motion racial/ethnic groups. The <u>standard</u> method corresponded to the brain-behavior associations derived from the full fMRI timeseries of the low-motion youth with a mean FD < 0.20 mm. The <u>motion-ordered</u> method corresponded to the brain-behavior associations derived from the scrubbed fMRI timeseries ranked and thresholded by their 100 least motion-corrupted timepoints (0 < FD < 0.20 mm) to construct the functional connectivity matrices of the low-motion youth. The <u>bagging</u> method corresponded to the brain-behavior associations derived from the scrubbed fMRI timeseries ranked and thresholded by their 100 least motion-corrupted timepoints (0 < FD < 0.20 mm) from which 100 timepoints were bootstrapped across 500 iterations to construct the functional connectivity matrices of the low-motion youth. The sample sizes were bootstrapped at 11 logarithmically-spaced *N* intervals: White *N* ∈ {25, 39, 62, 97, 152, 238, 374, 586, 920, 1444, 2266}; Black *N* ∈ {25, 33, 44, 58, 77, 101, 134, 177, 234, 310, 410}; Hispanic *N* ∈ {25, 35, 48, 67, 93, 129, 179, 249, 345, 480, 666}. For each low-motion racial/ethnic group and behavior, we computed a total of 11 intervals x 500 bootstrap *N* samples = 5,500 standard and motion-ordered correlations. For each low-motion racial/ethnic group and behavior, we computed a total of 11 intervals x 500 bootstrap *N* samples x 500 bootstrap timepoints = 2.75 million bagged correlations. For each racial/ethnic group, the brain-behavior *R*_s_ and 95% CI are shown as a function of sample size. The areas under the curve (AUC) for the brain-behavior associations also are displayed.

| ABCD Study | Brain-behavior relationships for NIH Toolbox |  |  |
| --- | --- | --- | --- |
|  | Standard | Motion-ordered | Bagging |
| **Low-motion White Youth** | **mean $R_s$ [95% CI]** | **mean $R_s$ [95% CI]** | **mean $R_s$ [95% CI]** |
| 25 | 0.10 [-0.32, 0.49] | 0.10 [-0.31, 0.50] | 0.11 [-0.28, 0.49] |
| 39 | 0.10 [-0.22, 0.42] | 0.088 [-0.23, 0.39] | 0.098 [-0.22, 0.43] |
| 62 | 0.097 [-0.14, 0.35] | 0.11 [-0.15, 0.34] | 0.094 [-0.17, 0.37] |
| 97 | 0.093 [-0.098, 0.29] | 0.11 [-0.077, 0.29] | 0.11 [-0.088, 0.28] |
| 152 | 0.11 [-0.057, 0.26] | 0.10 [-0.050, 0.26] | 0.11 [-0.038, 0.23] |
| 238 | 0.095 [-0.019, 0.21] | 0.11 [-0.011, 0.21] | 0.10 [-0.012, 0.23] |
| 374 | 0.10 [0.006, 0.19] | 0.099 [0.009, 0.20] | 0.11 [0.016, 0.19] |
| 586 | 0.10 [0.033, 0.18] | 0.11 [0.028, 0.17] | 0.11 [0.038, 0.17] |
| 920 | 0.10 [0.051, 0.15] | 0.11 [0.057, 0.15] | 0.11 [0.057, 0.15] |
| 1444 | 0.10 [0.073, 0.13] | 0.11 [0.074, 0.14] | 0.11 [0.076, 0.14] |
| 2266 | 0.10 [0.10, 0.10] | 0.11 [0.11, 0.11] | 0.11 [0.11, 0.11] |
| **Area Under Curve (AUC)** | **250.01** | **250.55** | **244.67** |
| **Low-motion Black Youth** | **mean $R_s$ [95% CI]** | **mean $R_s$ [95% CI]** | **mean $R_s$ [95% CI]** |
| 25 | 0.13 [-0.29, 0.51] | 0.12 [-0.23, 0.52] | 0.13 [-0.33, 0.50] |
| 33 | 0.12 [-0.22, 0.42] | 0.13 [-0.19, 0.44] | 0.11 [-0.22, 0.42] |
| 44 | 0.12 [-0.17, 0.39] | 0.14 [-0.14, 0.42] | 0.13 [-0.17, 0.42] |
| 58 | 0.11 [-0.12, 0.33] | 0.12 [-0.16, 0.38] | 0.14 [-0.092, 0.39] |
| 77 | 0.12 [-0.096, 0.30] | 0.13 [-0.078, 0.33] | 0.12 [-0.11, 0.34] |
| 101 | 0.12 [-0.060, 0.29] | 0.13 [-0.050, 0.28] | 0.12 [-0.050, 0.28] |
| 134 | 0.12 [-0.017, 0.25] | 0.13 [-0.005, 0.27] | 0.11 [-0.029, 0.24] |
| 177 | 0.12 [0.003, 0.23] | 0.13 [0.013, 0.24] | 0.12 [0.007, 0.22] |
| 234 | 0.12 [0.034, 0.21] | 0.12 [0.031, 0.21] | 0.12 [0.033, 0.20] |
| 310 | 0.12 [0.067, 0.18] | 0.13 [0.073, 0.18] | 0.12 [0.069, 0.18] |
| 410 | 0.12 [0.12, 0.12] | 0.13 [0.13, 0.13] | 0.12 [0.12, 0.12] |
| **Area Under Curve (AUC)** | **85.33** | **86.21** | **85.55** |
| **Low-motion Hispanic Youth** | **mean $R_s$ [95% CI]** | **mean $R_s$ [95% CI]** | **mean $R_s$ [95% CI]** |
| 25 | 0.049 [-0.34, 0.48] | 0.086 [-0.36, 0.51] | 0.062 [-0.39, 0.46] |
| 35 | 0.056 [-0.29, 0.38] | 0.063 [-0.27, 0.41] | 0.080 [-0.26, 0.40] |
| 48 | 0.056 [-0.24, 0.35] | 0.060 [-0.23, 0.33] | 0.064 [-0.22, 0.35] |
| 67 | 0.044 [-0.19, 0.25] | 0.058 [-0.14, 0.28] | 0.068 [-0.15, 0.29] |
| 93 | 0.056 [-0.14, 0.25] | 0.064 [-0.11, 0.25] | 0.064 [-0.11, 0.25] |
| 129 | 0.055 [-0.11, 0.21] | 0.066 [-0.091, 0.22] | 0.062 [-0.086, 0.21] |
| 179 | 0.052 [-0.065, 0.17] | 0.065 [-0.056, 0.18] | 0.064 [-0.066, 0.18] |
| 249 | 0.055 [-0.047, 0.15] | 0.066 [-0.032, 0.16] | 0.066 [-0.044, 0.16] |
| 345 | 0.054 [-0.035, 0.13] | 0.066 [-0.005, 0.15] | 0.064 [-0.010, 0.14] |
| 480 | 0.056 [0.004, 0.11] | 0.067 [0.021, 0.11] | 0.065 [0.018, 0.11] |
| 666 | 0.054 [0.054, 0.054] | 0.066 [0.066, 0.066] | 0.066 [0.064, 0.067] |
| **Area Under Curve (AUC)** | **123.47** | **117.35** | **118.78** |

**Table 2.** Correlations between functional connectivity and CBCL-externalizing obtained from the standard, motion-ordered, and bagging methods as a function of sample size across the 3 low-motion racial/ethnic groups. The <u>standard</u> method corresponded to the brain-behavior associations derived from the full fMRI timeseries of the low-motion youth with a mean FD < 0.20 mm. The <u>motion-ordered</u> method corresponded to the brain-behavior associations derived from the scrubbed fMRI timeseries ranked and thresholded by their 100 least motion-corrupted timepoints (0 < FD < 0.20 mm) to construct the functional connectivity matrices of the low-motion youth. The <u>bagging</u> method corresponded to the brain-behavior associations derived from the scrubbed fMRI timeseries ranked and thresholded by their 100 least motion-corrupted timepoints (0 < FD < 0.20 mm) from which 100 timepoints were bootstrapped across 500 iterations to construct the functional connectivity matrices of the low-motion youth. The sample sizes were bootstrapped at 11 logarithmically-spaced *N* intervals: White *N* ∈ {25, 39, 62, 97, 152, 238, 374, 586, 920, 1444, 2266}; Black *N* ∈ {25, 33, 44, 58, 77, 101, 134, 177, 234, 310, 410}; Hispanic *N* ∈ {25, 35, 48, 67, 93, 129, 179, 249, 345, 480, 666}. For each low-motion racial/ethnic group and behavior, we computed a total of 11 intervals x 500 bootstrap *N* samples = 5,500 standard and motion-ordered correlations. For each low-motion racial/ethnic group and behavior, we computed a total of 11 intervals x 500 bootstrap *N* samples x 500 bootstrap timepoints = 2.75 million bagged correlations. For each racial/ethnic group, the brain-behavior *R*_s_ and 95% CI are shown as a function of sample size. The areas under the curve (AUC) for the brain-behavior associations also are displayed.

| ABCD Study | Brain-behavior relationships for CBCL-externalizing |  |  |
| --- | --- | --- | --- |
|  | Standard | Motion-ordered | Bagging |
| **Low-motion White Youth** | **mean $R_s$ [95% CI]** | **mean $R_s$ [95% CI]** | **mean $R_s$ [95% CI]** |
| 25 | 0.097 [-0.29, 0.46] | 0.064 [-0.34, 0.49] | 0.071 [-0.32, 0.44] |
| 39 | 0.10 [-0.19, 0.41] | 0.086 [-0.22, 0.39] | 0.067 [-0.26, 0.40] |
| 62 | 0.094 [-0.13, 0.33] | 0.063 [-0.20, 0.30] | 0.067 [-0.20, 0.30] |
| 97 | 0.092 [-0.099, 0.28] | 0.066 [-0.12, 0.25] | 0.068 [-0.12, 0.27] |
| 152 | 0.098 [-0.050, 0.25] | 0.072 [-0.078, 0.22] | 0.066 [-0.082, 0.22] |
| 238 | 0.095 [-0.033, 0.22] | 0.069 [-0.049, 0.18] | 0.073 [-0.042, 0.19] |
| 374 | 0.099 [0.007, 0.19] | 0.072 [-0.027, 0.16] | 0.070 [-0.023, 0.16] |
| 586 | 0.10 [0.031, 0.17] | 0.072 [-0.006, 0.14] | 0.070 [0.005, 0.15] |
| 920 | 0.098 [0.053, 0.14] | 0.070 [0.022, 0.12] | 0.070 [0.019, 0.12] |
| 1444 | 0.099 [0.068, 0.13] | 0.071 [0.038, 0.10] | 0.072 [0.041, 0.10] |
| 2266 | 0.099 [0.099, 0.099] | 0.071 [0.071, 0.071] | 0.071 [0.071, 0.072] |
| **Area Under Curve (AUC)** | **244.82** | **250.70** | **251.44** |
| **Low-motion Black Youth** | **mean $R_s$ [95% CI]** | **mean $R_s$ [95% CI]** | **mean $R_s$ [95% CI]** |
| 25 | 0.055 [-0.35, 0.45] | 0.078 [-0.31, 0.48] | 0.039 [-0.36, 0.43] |
| 33 | 0.060 [-0.28, 0.38] | 0.063 [-0.30, 0.42] | 0.045 [-0.30, 0.39] |
| 44 | 0.056 [-0.23, 0.35] | 0.052 [-0.23, 0.36] | 0.050 [-0.22, 0.31] |
| 58 | 0.054 [-0.18, 0.27] | 0.056 [-0.18, 0.29] | 0.051 [-0.18, 0.28] |
| 77 | 0.048 [-0.15, 0.23] | 0.047 [-0.15, 0.26] | 0.046 [-0.16, 0.25] |
| 101 | 0.055 [-0.12, 0.22] | 0.055 [-0.10, 0.22] | 0.044 [-0.14, 0.22] |
| 134 | 0.061 [-0.065, 0.19] | 0.052 [-0.088, 0.17] | 0.053 [-0.094, 0.19] |
| 177 | 0.059 [-0.048, 0.16] | 0.055 [-0.055, 0.16] | 0.050 [-0.055, 0.17] |
| 234 | 0.063 [-0.023, 0.15] | 0.053 [-0.030, 0.14] | 0.053 [-0.026, 0.14] |
| 310 | 0.059 [0.007, 0.11] | 0.051 [-0.004, 0.11] | 0.052 [-0.005, 0.10] |
| 410 | 0.059 [0.059, 0.059] | 0.054 [0.054, 0.054] | 0.051 [0.049, 0.053] |
| **Area Under Curve (AUC)** | **82.69** | **84.96** | **85.52** |
| **Low-motion Hispanic Youth** | **mean $R_s$ [95% CI]** | **mean $R_s$ [95% CI]** | **mean $R_s$ [95% CI]** |
| 25 | 0.061 [-0.34, 0.43] | 0.065 [-0.33, 0.45] | 0.053 [-0.37, 0.45] |
| 35 | 0.079 [-0.22, 0.43] | 0.058 [-0.29, 0.38] | 0.065 [-0.25, 0.37] |
| 48 | 0.089 [-0.20, 0.34] | 0.066 [-0.18, 0.35] | 0.073 [-0.21, 0.35] |
| 67 | 0.081 [-0.15, 0.31] | 0.067 [-0.17, 0.31] | 0.073 [-0.16, 0.32] |
| 93 | 0.086 [-0.091, 0.27] | 0.065 [-0.11, 0.27] | 0.067 [-0.11, 0.26] |
| 129 | 0.083 [-0.063, 0.23] | 0.064 [-0.082, 0.22] | 0.065 [-0.085, 0.22] |
| 179 | 0.082 [-0.036, 0.21] | 0.063 [-0.056, 0.19] | 0.073 [-0.039, 0.19] |
| 249 | 0.077 [-0.016, 0.16] | 0.064 [-0.034, 0.15] | 0.069 [-0.015, 0.16] |
| 345 | 0.082 [0.007, 0.14] | 0.068 [-0.008, 0.14] | 0.068 [-0.008, 0.14] |
| 480 | 0.082 [0.037, 0.13] | 0.066 [0.021, 0.11] | 0.067 [0.018, 0.11] |
| 666 | 0.082 [0.082, 0.082] | 0.066 [0.066, 0.066] | 0.066 [0.065, 0.068] |
| **Area Under Curve (AUC)** | **113.75** | **117.00** | **115.94** |

### Motion-ordered and bagged brain-behavior associations from low-motion racial and ethnic groups

Following our previous work^29^, we next assessed the reproducibility of the motion-ordered and bagged brain-behavior relationships as a function of *N* ranging from typical (*N* = 25) to large (maximum *N* of each group). For the motion-ordered method, functional connectivity was computed with 100 least motion-corrupted timepoints in the scrubbed fMRI timeseries of the low-motion youth (**Online Methods**). For bagging, functional connectivity was computed with 100 least motion-corrupted timepoints in the scrubbed fMRI timeseries of the low-motion youth from which 100 timepoints were bootstrapped *with* replacement over 500 iterations (**Online Methods**). Similar to the standard method, we observed the presence of sampling variability due to the tightening of the 95% CI as the sample sizes of the racial/ethnic groups increased from typical to large for the NIH Toolbox (**Table 1**) and CBCL-externalizing (**Table 2**). The brain-behavior relationships were comparable to those obtained from the standard method. For NIH Toolbox, White 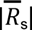 = 0.11, Black 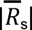 = 0.13, and Hispanic 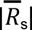 = 0.066 using the motion-ordered method, and White 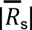 = 0.11, Black 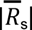 = 0.12, and Hispanic 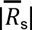 = 0.066 using bagging. For CBCL-externalizing, White 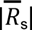 = 0.071, Black 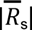 = 0.056, and Hispanic 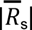 = 0.065 using the motion-ordered method, and White 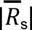 = 0.070, Black 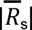 = 0.049, and Hispanic 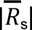 = 0.067 using bagging. To compare the performances of the motion-ordered and bagging methods with the standard method, we estimated the absolute areas under the curve (AUC) for each low-motion racial/ethnic group and behavior. Specifically, we computed the ΔAUC which quantifies the rate of change in AUC between the standard and motion-ordered as well as bagging methods. For NIH Toolbox, ΔAUC_motion-ordered_ was as follows: White ΔAUC = 0.22%, Black ΔAUC = 1.03%, and Hispanic ΔAUC = 4.96%. For NIH Toolbox, ΔAUC_bagging_ was as follows: White ΔAUC = 2.14%, Black ΔAUC = 0.26%, and Hispanic ΔAUC = 3.80%. For CBCL-externalizing, ΔAUC_motion-ordered_ was as follows: White ΔAUC = 2.40%, Black ΔAUC = 2.75%, and Hispanic ΔAUC = 2.86%. ForCBCL-externalizing, ΔAUC_bagging_ was as follows: White ΔAUC = 2.70%, Black ΔAUC = 3.42%, and Hispanic ΔAUC = 1.93%.

### Retaining high-motion youth for inclusive and reproducible brain-behavior associations

We next tested whether motion-ordered and bagging methods can maximize participant inclusion for generating inclusive and reproducible brain-behavior associations. Instead of discarding all the “high-motion” youth based on common practices in brain-behavior association studies, such as applying a mean FD < 0.20 mm cutoff, we aimed to retain all the low-/high-motion youth who had “usable” timepoints in their fMRI timeseries. To maximize participant inclusion, we retained all the low-/high-motion youth who had a minimum of 100 least motion-corrupted timepoints (minTP) in their scrubbed fMRI timeseries (**Online Methods**). This means that a given youth may have a mean FD > 0.20 mm (and would therefore normally be excluded), but, providing they have at least 100 (minTP) with FD < 0.20 mm timepoints in their motion-limited timeseries, they were included in analyses. This procedure helped to retain a total of 5,732 participants, corresponding to ∼99.98% of the total sample size: White *N* = 3,675, Black *N* = 810, and Hispanic *N* = 1,247. Fundamentally, this procedure maximized the representation of minoritized youth for the subsequent brain-behavior analyses: Loss in White *N*: 0.03%, Loss in Black *N*: 0%, and Loss in Hispanic *N*: 0%.

We reexamined the reproducibility of the motion-ordered and bagged brain-behavior associations as a function of *N* ranging from typical (*N* = 25) to large (maximum *N* of each group). We computed the standard brain-behavior associations to compare the performances of the motion-ordered and bagging methods when the high-motion youth were included. We observed similar narrowing of the 95% CIs with the standard and motion-ordered methods (**Fig.3**) in addition to the standard and bagging methods (**Fig.4**) as the sample sizes of the racial/ethnic groups grew from typical to large for the NIH Toolbox and CBCL-externalizing. The standard, motion-ordered, and bagged brain-behavior associations remained comparable even when the “high-motion” youth were included. From the standard method and NIH Toolbox, White 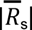 = 0.090, Black 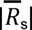 = 0.076, and Hispanic 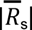 = 0.047. From the motion-ordered method and NIH Toolbox, White 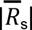 = 0.086, Black 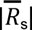 = 0.071, and Hispanic 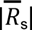 = 0.044. From bagging and NIH Toolbox, White 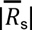 = 0.085, Black 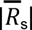 = 0.072, and Hispanic 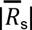 = 0.043. From the standard method and CBCL-externalizing, White 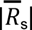 = 0.065, Black 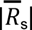 = 0.069, and Hispanic 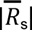 = 0.034. From the motion-ordered method and CBCL-externalizing, White 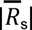 = 0.056, Black 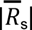 = 0.049, and Hispanic 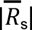 = 0.037. From bagging and CBCL-externalizing, White 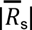 = 0.055, Black 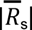 = 0.045, and Hispanic 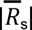 = 0.041. The absolute AUC_motion-ordered_ for NIH Toolbox were as follows: White ΔAUC = 0.72%, Black ΔAUC = 2.58%, and Hispanic ΔAUC = 0.09%. For NIH Toolbox, ΔAUC_bagging_ was as follows: White ΔAUC = 0.55%, Black ΔAUC = 3.03%, and Hispanic ΔAUC = 3.87%. For CBCL-externalizing, Δ AUC_motion-ordered_ was as follows: White ΔAUC = 3.05%, Black ΔAUC = 2.15%, and Hispanic ΔAUC = 3.03%. For CBCL-externalizing, ΔAUC_bagging_ was as follows: White ΔAUC = 0.37%, Black ΔAUC = 2.00%, and Hispanic ΔAUC = 1.47%.

**Fig. 3.**
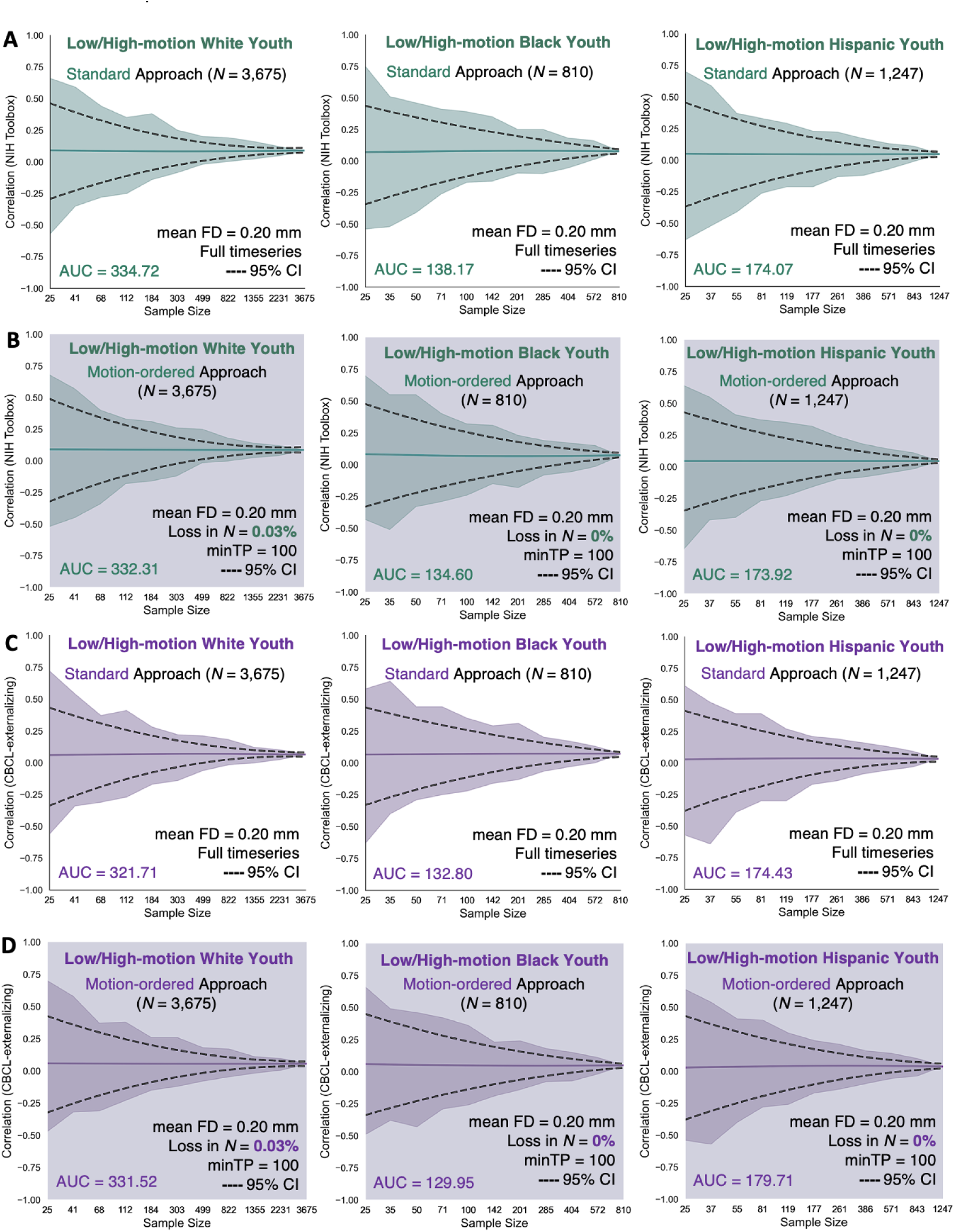
Correlations between functional connectivity and (A-B) NIH Toolbox and (C-D) CBCL-externalizing obtained from the standard and motion-ordered methods as a function of sample size across the 3 low-/high-motion racial/ethnic groups. The <u>standard method</u> corresponded to the brain-behavior associations derived from the full fMRI timeseries of the low-/high-motion youth that have been retained for the analyses without imposing an initial head motion threshold of mean FD < 0.20 mm. The <u>motion-ordered method</u> corresponded to the brain-behavior associations derived from the scrubbed fMRI timeseries of the low-/high-motion youth who were retained based on the assumption that they had a minimum of 100 least motion-corrupted timepoints. Their scrubbed fMRI timeseries were ranked by their lowest FD values and 100 least motion-corrupted timepoints (0 < FD < 0.20 mm) were selected to construct the functional connectivity matrices of the youth. The sample sizes were bootstrapped at 11 logarithmically-spaced *N* intervals: White *N* ∈ {25, 41, 68, 112, 184, 303, 499, 822, 1355, 2231, 3675}; Black *N* ∈ {25, 35, 50, 71, 100, 142, 201, 285, 404, 572, 810}; Hispanic *N* ∈ {25, 37, 55, 81, 119, 177, 261, 386, 571, 843, 1247}. For each low-/high-motion racial/ethnic group and behavior, we computed a total of 11 intervals x 500 bootstrap *N* samples = 5,500 standard and motion-ordered correlations. Solid teal and purple lines show the mean correlations from the 500 bootstrap samples for a given same size. Teal and purple shadings denote the minimum and maximum correlations across 500 bootstrap samples for a given sample size. Black dotted lines represent the lower and upper bounds of the 95% CIs for a given sample size. The areas under the curve (AUC) for the brain-behavior associations also are displayed.

**Fig. 4.**
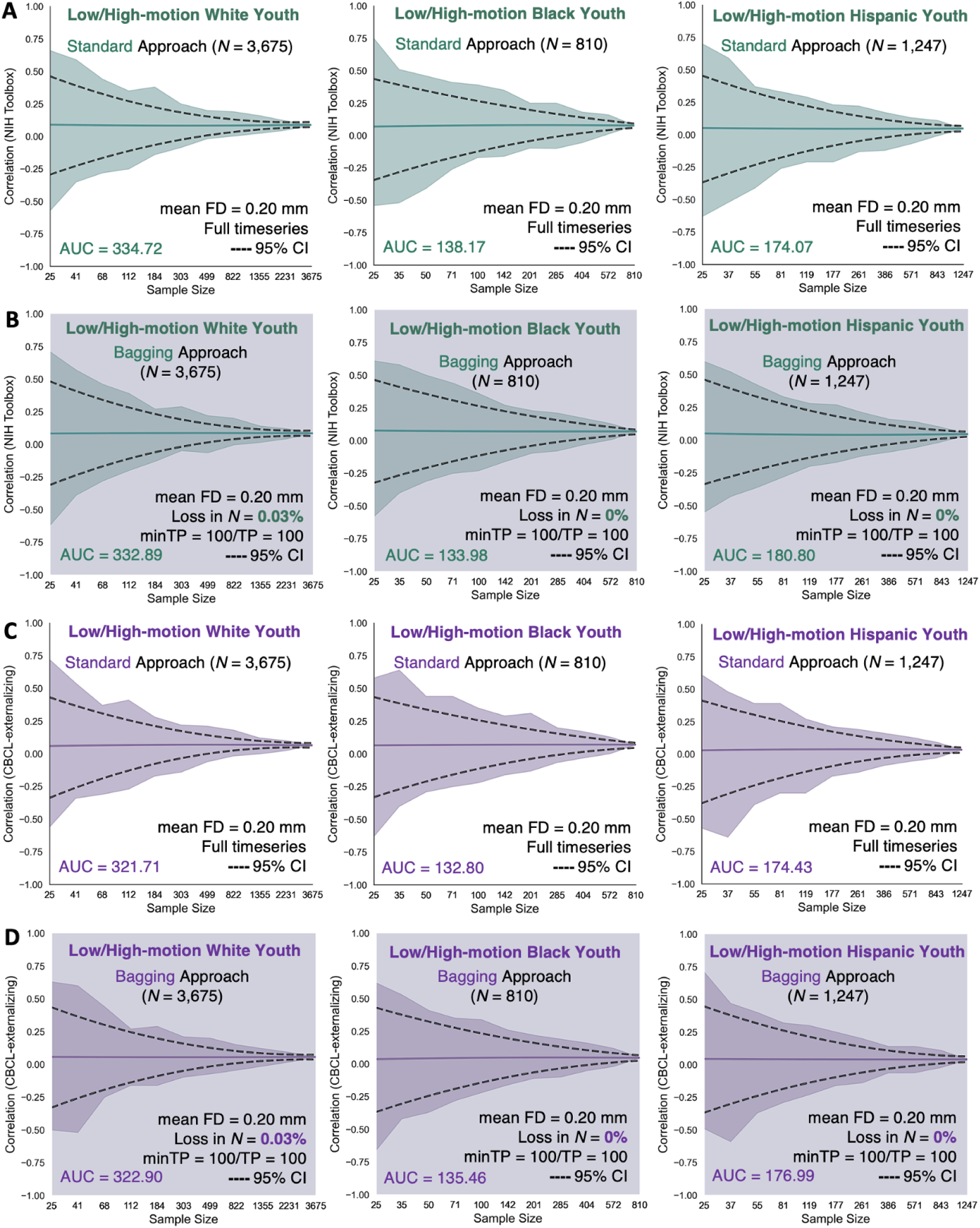
Correlations between functional connectivity and (A-B) NIH Toolbox and (C-D) CBCL-externalizing obtained from the standard and bagging methods as a function of sample size across the 3 low-/high-motion racial/ethnic groups. The <u>standard method</u> corresponded to the brain-behavior associations derived from the full fMRI timeseries of the low-/high-motion youth that have been retained for the analyses without imposing an initial head motion threshold of mean FD < 0.20 mm. The <u>bagging method</u> corresponded to the brain-behavior associations derived from the scrubbed fMRI timeseries of the low-/high-motion youth who were retained based on the assumption that they had a minimum of 100 least motion-corrupted timepoints. Their scrubbed fMRI timeseries were ranked by their lowest FD values and 100 least motion-corrupted timepoints (0 < FD < 0.20 mm) were selected from which 100 timepoints were bootstrapped across 500 iterations to construct the functional connectivity matrices of the youth. The sample sizes were bootstrapped at 11 logarithmically-spaced *N* intervals: White *N* ∈ {25, 41, 68, 112, 184, 303, 499, 822, 1355, 2231, 3675}; Black *N* ∈ {25, 35, 50, 71, 100, 142, 201, 285, 404, 572, 810}; Hispanic *N* ∈ {25, 37, 55, 81, 119, 177, 261, 386, 571, 843, 1247}. For each low-/high-motion racial/ethnic group and behavior, we computed a total of 11 intervals x 500 bootstrap *N* samples = 5,500 standard correlations. For each low-/high-motion racial/ethnic group and behavior, we computed a total of 11 intervals x 500 bootstrap *N* samples x 500 bootstrap timepoints = 2.75 million bagged correlations. Solid teal and purple lines show the mean correlations from the 500 bootstrap samples for a given same size. Teal and purple shadings denote the minimum and maximum correlations across 500 bootstrap samples for a given sample size. Black dotted lines represent the lower and upper bounds of the 95% CIs for a given sample size. The areas under the curve (AUC) for the brain-behavior associations also are displayed.

### Concordance of standard and motion-ordered/bagged brain-behavior relationships from low-/high-motion youth

We evaluated the concordance of the standard and motion-ordered/bagged brain-behavior relationships using the Lin’s concordance correlation coefficient (ρ_c_)^37,38^ across the 3 racial/ethnic groups. By definition, ρ_c_ measures the agreement between two variables from a set of bivariate data (**Online Methods**). It can be used to quantify the degree to which one measurement reproduces another measurement. ρ_c_ can be seen as a weighted version of the Pearson correlation coefficient (*r*) ranging from −1 to 1. We also computed *r* between the standard and motion-ordered/bagged brain-behavior relationships to show that these two metrics are not identical. We derived ρ_c_ and *r* from the low-motion youth at typical (*N* = 25) and large (*N* = 250) sample sizes. We then derived ρ_c_ and *r* from the low-/high-motion youth who were retained based on their 100 least motion-corrupted timepoints at typical (*N* = 25) and large (*N* = 250) sample sizes. At typical sample sizes, we observed strong concordance between the standard and motion-ordered brain-behavior relationships for the NIH Toolbox (0.76 < ρ_c_ < 0.83) and CBCL-externalizing (0.77 < ρ_c_ < 0.83) derived from the low-motion racial/ethnic groups (**Fig.5A&B**). Similar concordance patterns were observed at typical sample sizes when the high-motion youth were included for the NIH Toolbox (0.76 < ρ_c_ < 0.79) and CBCL-externalizing (0.69 < ρ_c_ < 0.80) (**Fig.5C&D**). At larger sample sizes, the concordance between the standard and motion-ordered brain-behavior relationships remained strong for the NIH Toolbox (0.78 < ρ_c_ < 0.81) and CBCL-externalizing (0.76 < ρ_c_ < 0.83) obtained from the low-motion racial/ethnic groups (**Fig.6A&B**). When the high-motion youth were included at larger sample sizes, the concordance persisted for the NIH Toolbox (0.79 < ρ_c_ < 0.82) and CBCL-externalizing (0.72 < ρ_c_ < 0.80) (**Fig.6C&D**). Moreover, *r* consistently was strong across typical and large sample sizes for the two behaviors both when high-motion youth were included *and* when they were excluded (**Fig.5&6**). Additionally, the concordance and similarity of the standard and bagged brain-behavior associations were consistent with those derived from the motion-ordered method at typical and large sample sizes (**Supplementary Figures 3-4**).

**Fig. 5.**
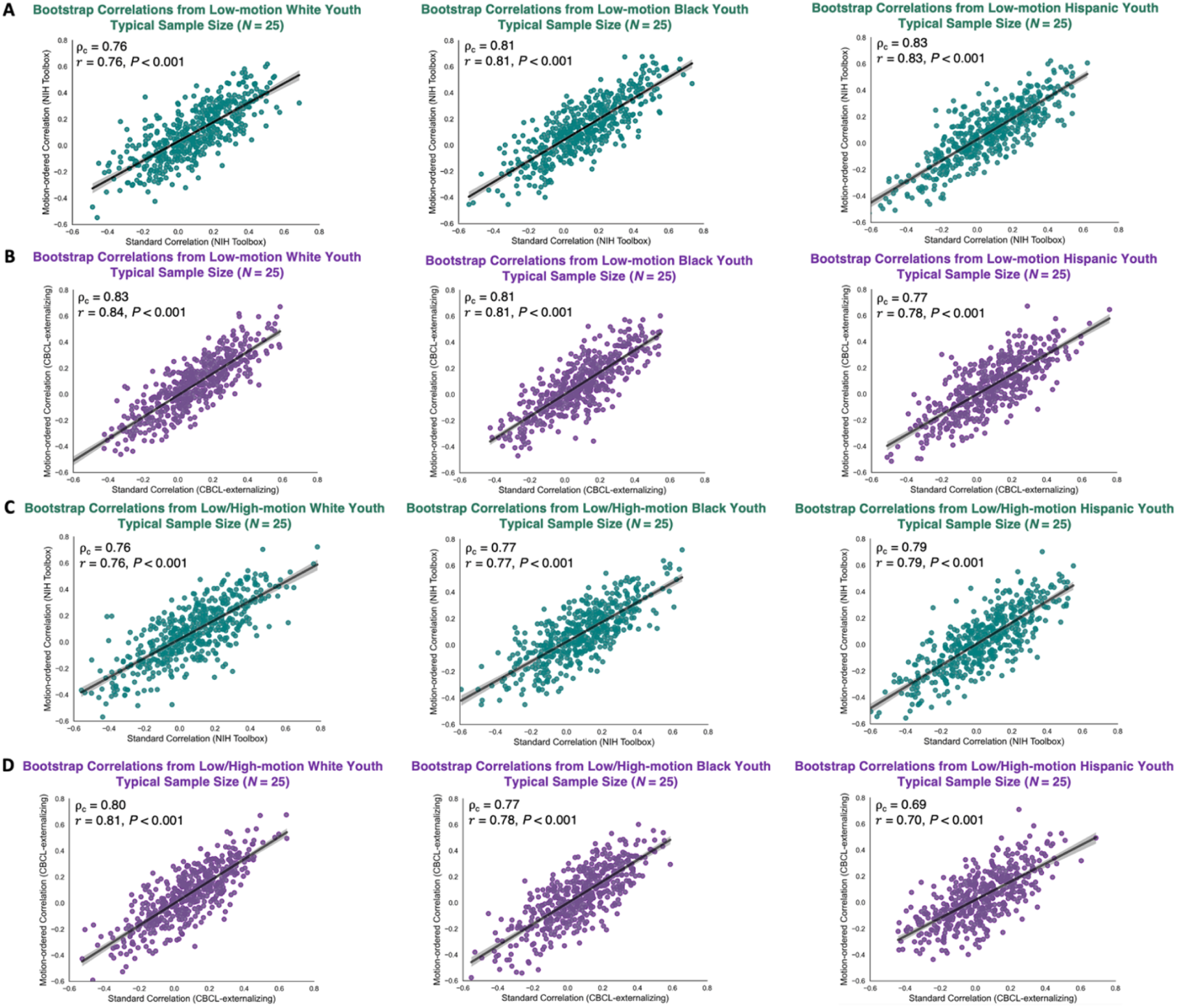
Bootstrap brain-behavior correlations obtained from the standard and motion-ordered methods at typical sample size (*N* = 25) *with* (A-B) and *without* (C-D) excluding the high-motion youth across the 3 racial/ethnic groups. The teal and purple data points corresponded to bootstrap samples at *N* = 25 obtained from 500 iterations and these samples were used to compute the associations between functional connectivity and NIH Toolbox and CBCL-externalizing. The Lin’s concordance correlation coefficient (ρ_c_) and Pearson correlation coefficient (*r*) were computed respectively to assess the reproducibility and similarity of the brain-behavior associations obtained from the standard and motion-ordered methods. The low-/high-motion youth were retained based on the assumption that they had a minimum of 100 least motion-corrupted timepoints in their fMRI timeseries.

**Fig. 6.**
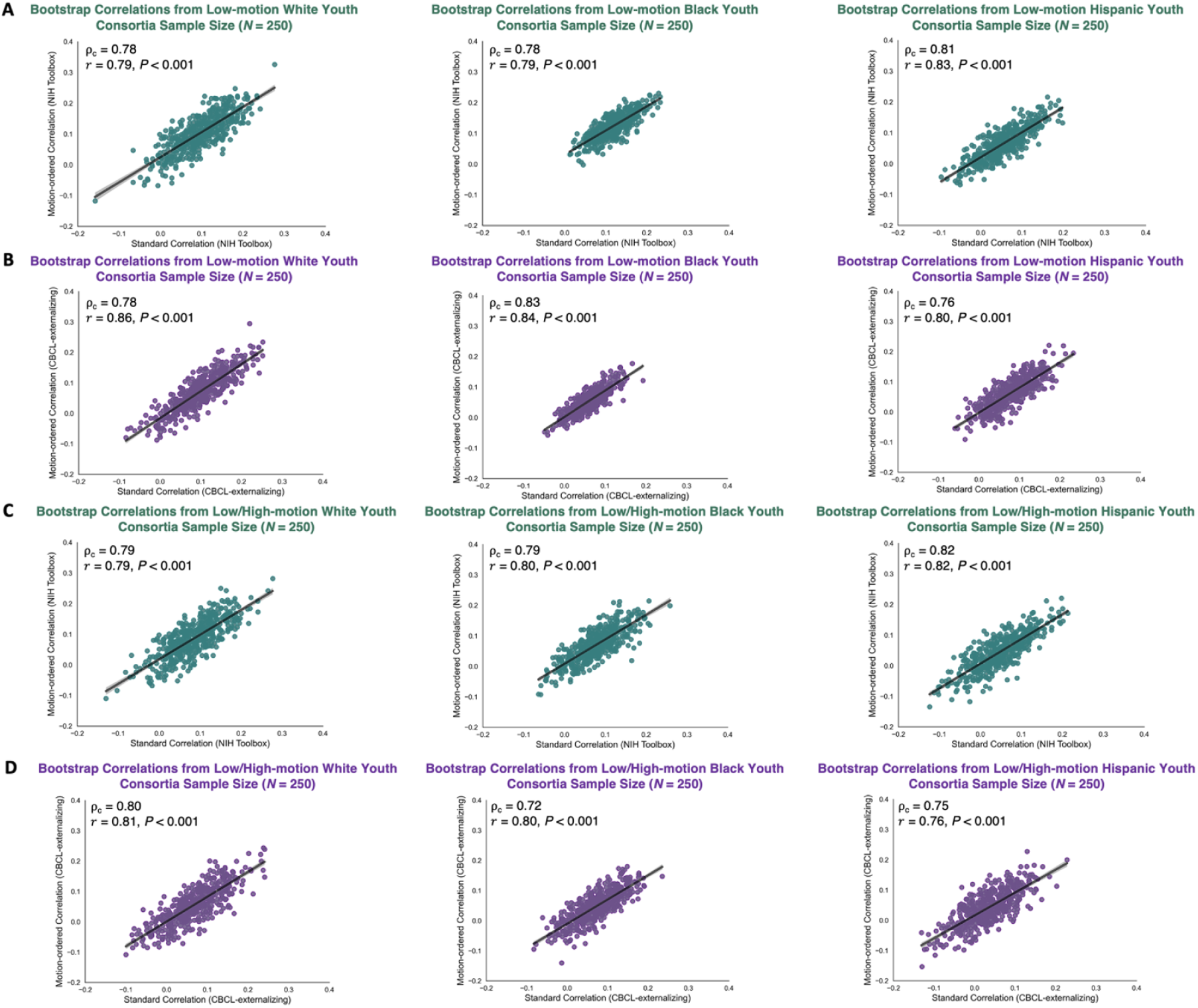
Bootstrap brain-behavior correlations obtained from the standard and motion-ordered methods at large sample size (*N* = 250) *with* (A-B) and *without* (C-D) excluding the high-motion youth across the 3 racial/ethnic groups. The teal and purple data points corresponded to bootstrap samples at *N* = 250 obtained from 500 iterations and these samples were used to compute the associations between functional connectivity and NIH Toolbox and CBCL-externalizing. The Lin’s concordance correlation coefficient (ρ_c_) and Pearson correlation coefficient (*r*) were computed respectively to assess the reproducibility and similarity of the brain-behavior associations obtained from the standard and motion-ordered methods. The low-/high-motion youth were retained based on the assumption that they had a minimum of 100 least motion-corrupted timepoints in their fMRI timeseries.

The standard, motion-ordered, and bagged brain-behavior relationships were computed by estimating functional connectivity from the fMRI timeseries of the youth that were concatenated across a maximum of 4 resting-state fMRI scans. We tested whether the motion-ordered and bagging methods could still maximize the representation of the minoritized participants to produce inclusive and reproducible brain-behavior relationships using only one fMRI scan (REST1) lasting for 5 min. This is because community-based samples and consortia-level datasets do not necessarily acquire repeated scans within- or between-sessions^39^. From REST1, a total of 4,736 low-motion youth were identified with a mean FD < 0.20 mm, representing ∼83% of the total sample size (**Supplementary Figure 5**): White *N* = 3,140, Black *N* = 600, and Hispanic *N* = 996. To maximize participant inclusion, we observed that the motion-ordered and bagging methods could be performed using 100 least motion-corrupted timepoints from the REST1 scrubbed timeseries (**Supplementary Figure 5**). For comparison purposes, we retained low-motion youth who had a minimum of 100 least motion-corrupted timepoints from REST1: White *N* = 3,120, Black *N* = 596, and Hispanic *N* = 990. Subsequently, we retained all the low-/high-motion youth who had a minimum of 100 least motion-corrupted timepoints from REST1. This procedure increased the sample size to 5,526 youth which corresponded to ∼96% of the total sample size. The loss in *N* across the 3 racial/ethnic groups was as follows: Loss in White *N* = 2.9%, Loss in Black *N* = 5.8%, and Loss in Hispanic *N* = 4.4%. It is possible to reduce the minimum number of sampled timepoints to 50 as REST1 had considerably fewer timepoints compared to the concatenated timeseries across a maximum of 4 runs. This further reduced the loss in *N* across the 3 racial/ethnic groups: Loss in White *N* = 0.7%, Loss in Black *N* = 2.0%, and Loss in Hispanic *N* = 1.3%. However, determining the minimum number of sampled timepoints exceeds the scope of this study and is not practical as we cannot recommend a one-size-fits-all number for all typically sized and consortia datasets. All the effect sizes produced by the standard, motion-ordered, and bagging methods were comparable for the NIH Toolbox and CBCL-externalizing across the 3 racial/ethnic groups (**Supplementary Figures 6-9**).

The standard, motion-ordered, and bagged brain-behavior associations were computed using the edge that produced the strongest *R*_s_ in the functional connectome after surviving the Benjamini-Hochberg False Discovery Rate (q = 0.05)^36^. The strongest edges respectively had a correlation strength of |*R*_s_| = 0.20 for the NIH Toolbox and |*R*_s_| = 0.093 for the CBCL-externalizing. An edge may exhibit high (low) test-retest reliability but demonstrate low (high) behavioral relevance^40^. Therefore, we recomputed the standard, motion-ordered, and bagged brain-behavior associations using the weakest edge that shared a significant association with NIH Toolbox (|*R*_s_| = 0.042) and CBCL-externalizing (|*R*_s_| = 0.070) with and without including the high-motion youth who had a minimum of 100 least motion-corrupted timepoints in their fMRI timeseries. All the effect sizes produced by the standard, motion-ordered, and bagging methods were comparable for the NIH Toolbox and CBCL-externalizing across the 3 racial/ethnic groups (**Supplementary Figures 10-13**).

## Discussion

In modern neuroimaging research, the quest for reproducible brain-behavior relationships has underscored a need for large sample sizes that can provide adequate statistical power. At the same time, consortia datasets tend to undergo rigorous data quality control to mitigate the impact of noise, such as that produced by participant head motion, with strict thresholds for data inclusion. This creates a tension: applying strict head motion thresholds drastically reduces sample size, particularly in minoritized developmental cohorts. Here, replicating and extending previous work^29^, we demonstrated the utility of motion-ordering and motion-ordering+resampling (i.e., bagging) to preserve fMRI datasets containing high levels of head motion. Although minoritized youth exhibited greater head motion relative to White youth, both methods retained more than 99% of all minoritized youth and produced inclusive and reproducible effect sizes across races/ethnicities based on their motion-limited fMRI data. Overall, the motion-ordered and bagging methods enhance sample representation in brain-behavior association studies and fulfill the promise of consortia datasets to produce generalizable effect sizes across sociodemographically diverse cohorts, however, motion-ordering may be preferred for brain-behavior association studies due to its considerably lower computational demands relative to bagging^29^.

We showed that fMRI datasets containing head motion can be retained and used to compute meaningful brain-behavior associations for general cognitive performance and externalizing psychopathology. Applying the motion-ordered and bagging methods produced comparable, if not stronger, brain-behavior correlations compared with the standard motion correction method when high-motion datasets collected from minoritized youth were retained. The effect sizes from the standard versus motion-ordered and bagging methods differed by a maximum of 7%. Some may consider this a small gain, but it is important to note that the proposed methods effect sizes were obtained by including more than 99% of White, Black, and Hispanic youth. This held true when using a single fMRI scan and the weakest edge in the functional connectome. This extends previous work^29^ showing that we may not need to discard thousands of participants in consortia datasets, particularly across minoritized individuals, to produce meaningful brain-behavior associations. Individuals do not need to be excluded based on measures of framewise displacement per volume, but rather that they can be retained if they have enough motion-limited timeseries data.

We focused on generating inclusive and reproducible brain-behavior relationships by retaining a large proportion of data collected from minoritized youth. Increasingly, researchers also are interested in using brain-based models to predict behavior. However, these predictive models often are trained on samples that lack diversity, resulting in poorer prediction of behavior in minoritized individuals. For example, models trained on resting-state functional connectivity data have been found to predict a broad range of behaviors more accurately in White Americans relative to African Americans. Cognitive scores were found to be underpredicted in African Americans compared to White Americans in the ABCD Study, though predictive performance of the minoritized youth improved when the models were trained only on African Americans^41^. In contrast, externalizing and internalizing behaviors were overpredicted in African Americans compared with White Americans^41^. Brain-behavior models also can fail to predict certain behaviors from those who do not fit common neurocognitive profiles^42^. Not only has model failure — misclassification of neurocognitive scores as low or high from functional connectivity — been shown to be generalizable across datasets, behaviors, and participant groups, it can be associated with sociodemographic variables^42^. For example, high-scoring participants who belonged to non-White groups have been shown to be misclassified frequently as low-scoring and vice-versa for the White (not Hispanic/Latino) participants^42^. Ultimately, before moving to prediction, it is important for researchers to enhance sample representation so that the data used to train these predictive models is inclusive. The motion-ordering and bagging methods provide a way for improving the retention of minoritized individuals and securing a diverse sample.

Before concluding, it is important to note some limitations of the motion-ordered and bagging methods. First, there is no one-size-fits-all number of timepoints that can be recommended to generate inclusive and reproducible brain-behavior relationships. This is because typical samples and consortia datasets tend to acquire different number of frames as a result of varying scan lengths and sequence parameters. Still, we achieved comparable brain-behavior associations from both single and repeated fMRI scans. The focus should be on obtaining a set of least motion-corrupted timepoints from which data can be resampled, which is of particular importance for developmental and clinical cohorts, as those participants who are excluded for motion artifacts are usually those who are the youngest^13,21,22^ or who exhibit more severe symptoms/difficulties (e.g., ADHD, ASD)^43,44^. Retaining their data would not only maximize inclusivity but also enable a wider spectrum of behaviors to be investigated. Second, the proposed methods are effective for functional connectivity applications that do not rely on the continuity of the fMRI timeseries. The ranking of the motion-limited timepoints may not be desirable for traditional general linear models and intersubject correlations as they respectively require *a priori* information about the timecourse of the stimulus and contiguous timepoints to measure brain synchrony across individuals^45^. Third, the two methods require the fMRI timeseries to be scrubbed using a head motion threshold. If a study acquires a small number of timepoints, then scrubbing may still exclude a large number of participants due to insufficient amount of motion-limited timepoints. In the present study, one White high-motion youth could not be salvaged due to the threshold of minTP = 100 although all the Black and Hispanic high-motion youth were retained. Therefore, it is not practical to assume that the motion-ordered and bagging methods will always guarantee the retention of all the high-motion participants. Finally, the motion-ordered and bagging methods were developed to compute brain-behavior associations at the edge-level as a function of sample size. Given that edges in the functional connectome share variable behavioral relevance^40^, some may not exhibit any meaningful relationship with a behavioral domain. In the present study, there was no edge in the functional connectome that shared a significant association with internalizing behaviors. Future work could expand the proposed methods to examine their abilities in producing inclusive and reproducible brain-behavior associations at different levels of functional organization (e.g., component, network)^1^.

In conclusion, we can generate inclusive and reproducible brain-behavior relationships using motion-limited fMRI data from *low*- and *high*-motion youth across races and ethnicities. By maximizing participant inclusivity, motion-ordering affords a promising opportunity to retain high-motion youth, particularly those who belong to minoritized backgrounds, who otherwise would be discarded in typical brain-behavior association studies. It is our hope that motion-ordering will reduce the tension between data quality standards and sample size requirements, and in doing so, mitigate against resource wastage and the unethical practice of discarding large numbers of participants who have given their time, effort, and trust to the research process.

## Online Methods

### Participants

The Adolescent Brain Cognitive Development^SM^ Study (ABCD Study^®^) is a ten-year longitudinal and deeply phenotyped consortium that tracks the development of children and adolescents across 21 research sites in the United States^30^. The primary aim of the ABCD Study is to investigate how biological and environmental factors influence brain development as well as social and behavioral phenotypes over time^30,46,47^. In this study, we used the imaging, behavioral, and demographic data from the ABCD Study NIMH Data Archive Baseline Release 4.0 (*N* = 11,874; 9-10 years) (https://abcdstudy.org/). The youth were recruited from elementary schools based on their gender, race, ethnicity, socioeconomic status, and urbanicity^12^. The ABCD Study obtained approval from a centralized Institutional Review Board (IRB) located at the University of California, San Diego in addition to obtaining local IRB approval from each of the imaging sites^48^. Written assent was provided by the youth and written informed consent was obtained by their parents or guardians.

In this study, we included youth who belonged to White, Black, and Hispanic racial/ethnic groups. Youth who belonged to other races and ethnicities such as Asian, Mixed Race, and Native Americans were not included due to their limited sample sizes (∼13% of the total ABCD Study sample size). Only participants who had complete imaging, behavioral, and demographic data in addition to passing successfully the MRI data quality control criteria described by the ABCD Data Analysis and Informatics Center^49^ (DAIC; https://wiki.abcdstudy.org/release-notes/imaging/quality-control.html; imgincl_t1w_include = 1; imgincl_rsfmri_include = 1) were included in the study. The DAIC recommendations minimize data quality control failure from biasing the brain-behavior associations estimated from the low-/high-motion participants. A total of 5,733 youth remained in the study: White *N* = 3,676, Black *N* = 810, and Hispanic *N* = 1,247. The complete demographic information is presented in **Supplementary Table 1**.

### Behavioral Data

We focused on three measures related to cognitive performance, externalizing psychopathology, and internalizing psychopathology. The NIH Toolbox® (nihtoolbox.org/) was used to measure general cognitive performance based on the fluid and crystallized cognition composite scores of the youth^50–52^. For each youth, the total composite score of the NIH Toolbox was obtained which was *T*-standardized. The higher the NIH Toolbox *T*-score, the greater the level of cognitive functioning.

The Child Behavior Checklist (CBCL) is a parent report assessment that was used to measure externalizing and internalizing behaviors^53^. While externalizing behaviors tend to reflect conditions such as ADHD and oppositionality (e.g., impulsivity, aggression, disruption), internalizing behaviors capture mood-related behaviors (e.g., anxiety, withdrawal, dysphoria)^54^. The CBCL externalizing and internalizing scores of the youth were obtained from their respective syndrome scales that were subsequently *T*-standardized. The higher the CBCL externalizing and internalizing *T*-scores, the greater the risk of experiencing behavioral and emotional problems. On average, the youth exhibited weaker externalizing behaviors relative to internalizing behaviors in the baseline ABCD Study sample.

In youth, the test-retest reliability indexed by the intraclass correlation coefficient (ICC) is excellent for NIH Toolbox (ICC > 0.76)^55^ as well as CBCL externalizing and internalizing (ICC > 0.95)^53^. Similarly, the internal consistency measured by the Cronbach’s α is highly acceptable for NIH Toolbox (α > 0.77)^56^ as well as CBCL externalizing and internalizing (α > 0.90)^53^. Of note, CBCL-externalizing produced an inverse relationship with the NIH Toolbox (Externalizing: Spearman’s *R*_s_ = −0.11, *P* < 0.001; Internalizing: Spearman’s *R*_s_ = −0.005, *P* > 0.05). This relationship persisted after adjusting for baseline age, imaging site, area deprivation index, and pubertal status (**Supplementary Figure 14**). The behavioral information is presented in **Supplementary Table 1**.

### Consideration of Covariates

#### Head Motion

Head motion inflates functional connectivity estimates across different populations and it has an inverse relationship with age such that children tend to exhibit greater in-scanner movement relative to adults^13–20^. Although motion correction procedures are implemented during the pre-/post-processing stages, the influence of head motion on functional connectivity cannot be removed completely. Therefore, to minimize biases associated with head motion and spurious brain-behavior relationships in the youth sample, head motion (indexed by the mean FD) was treated as a covariate when computing the associations between functional connectivity-cognitive performance and connectivity-psychopathology (see **Head Motion** below).

#### Participant Sex Assigned at Birth

Sex assigned at birth differences are evident during brain development. The unique properties of an individual’s functional connectome (i.e., connectome distinctiveness) appear to be delayed in male compared to female youths^57^. Further, connectome-based predictions of cognitive functions differ in males and females based on their underlying within- and between-network connections^58^. Therefore, we treated participant sex assigned at birth^59^ as a covariate when computing the brain-behavior associations.

#### Baseline Age

In the present study, we used the ABCD Study NIMH Data Archive Baseline Release by recruiting youth aged from 9 to 10 years (see **Participants**). Due to the limited range of the baseline age, we did not treat age as a covariate when computing the brain-behavior associations. The age-adjusted edgewise correlations remained highly correlated with those obtained from accounting only for sex assigned at birth and head motion (**Supplementary Figure 15**).

#### Imaging Sites

The imaging sites of the youth were obtained for their baseline MRI scans across the United States^12^. While 21 sites actively acquire the imaging data for the youth over a ten-year period, the fMRI scanning protocols are optimized and harmonized across scanner manufacturers and sites (see **MRI Data Acquisition** below). Due to the harmonization sequence protocols that are in place to acquire the resting-state and task-based fMRI data, we did not adjust for imaging sites when computing the brain-behavior associations. The site-adjusted edgewise correlations also were highly correlated with those obtained from accounting only for sex assigned at birth and head motion (**Supplementary Figure 15**).

#### Area Deprivation Index

In the United States, unfortunately race often serves as a proxy for the structural and social disparities that exist in this country. Given the structural and social disparities that are present across races and ethnicities in the United States^60–62^, we considered the area deprivation index (ADI)^61,62^ which measures the socioeconomic status of the neighborhood they reside in. The ADI reflects the weighted sum of 17 composite scores relating to employment, education, income and poverty, and housing obtained at the census-tract level based on the youth’s home address. In our ABCD Study sample, we did not observe a significant difference in ADI across the 3 racial/ethnic groups: Kruskal-Wallis test, *H* (2) = 0.47, *P* = 0.79; White median ADI = 98.6, Black median ADI = 98.9, and Hispanic median ADI: 98.8. As the ADI-adjusted edgewise correlations remained highly correlated with those obtained from accounting only for sex assigned at birth and head motion (**Supplementary Figure 15**), ADI was not treated as a covariate when computing the brain-behavior associations.

#### Pubertal Status

The onset of puberty can affect brain development with hormonal changes contributing to physical, emotional, cognitive, and social alterations during adolescence^47^. We obtained the pubertal status of the youth from the male and female category subscales of the parent-report Pubertal Development Scale based on their sex assigned at birth^47,63^. As the puberty-adjusted edgewise correlations remained highly correlated with those obtained from accounting only for sex assigned at birth and head motion (**Supplementary Figure 15**), pubertal status was not treated as a covariate when computing the brain-behavior associations.

It is important to remember that controlling for covariates typically imposes a quandary in brain-behavior association studies since adjusting for too many or least potential confounds can impact reproducibility^64^.

### MRI Data Acquisition

Structural T1-weighted, T2-weighted, and functional MRI scans were acquired and optimized across 21 imaging sites in the United States using Siemens Prisma, Philips, and GE 750 3T scanners^30^. Two 3D MPRAGE T1-weighted and two 3D FSE T2-weighted volumes with spatial resolution 1 x 1 x 1 mm^3^ were obtained for each youth. A maximum of four runs of resting-state fMRI data were acquired and harmonized with a gradient-echo echo-planar imaging (EPI) sequence (60 slices, voxel size = 2.4 x 2.4 x 2.4 mm^3^, multiband factor = 6, TR = 800 ms, TE = 30 ms, flip angle = 52°). Each youth underwent a maximum of 20 min (4 runs x 5 min) resting-state scans while they fixated a cross on the screen. Real-time motion detection and correction configurations were set up for the structural scans using prospective motion correction (PROMO)^65^ on the GE scanners, and volumetric navigators (vNav)^66^ on the Siemens and Philips platforms. An fMRI integrated real-time motion monitoring system (FIRMM)^67^ also was implemented for the Siemens scanners to readjust the scanning paradigm based on the degree of head motion. The resting-state fMRI characteristics are presented in **Supplementary Table 1**.

### ABCD-BIDS Pipeline

All the ABCD MRI data were processed with the ABCD-BIDS Community Collection^31^ (ABCD-3165; https://github.com/DCAN-Labs/abcd-hcp-pipeline). The ABCD-BIDS pipeline is a modification of the HCP MRI pipeline^68^. This processing pipeline comprises five stages. (1) PreFreesurfer for performing brain extraction, denoising with ANTs^69^, and N4 bias field correction on the T1-weighted and/or T2-weighted scans. (2) FreeSurfer^70^ for constructing cortical surfaces from the normalized anatomical data. Anatomical segmentation, reconstruction of white/gray matter and gray matter/pial cortical surface boundaries, and surface registration to a standard surface template are performed. The surfaces are enhanced using the T2-weighted scans upon availability. The mid-surface thickness surfaces also are generated to capture the average white/gray matter and gray matter/CSF surfaces. (3) PostFreeSurfer for converting the PreFreesurfer and FreeSurfer outputs into the standard CIFTI template space. The volumes are transformed to the standard MNI space using ANTs non-linear registration. The surfaces also are transformed to the standard surface space from spherical registration using the ANTs deformable SyN algorithm. (4) fMRIVolume for registering the functional data to the standard MNI template. Each volume of the timeseries is aligned using FSL FLIRT^71^ rigid body registration to the initial frame to correct for motion. A pair of spin-echo EPI scans with reverse phase encoding directions are used for distortion corrections in the phase encoding direction of each fMRI volume using FSL’s toptup^71^. The single-band reference is then registered to the T1-weighted scans and this registration is used to align all the fMRI volumes to the anatomical data. Each fMRI volume is non-linearly registered to the standard MNI template using FSL FNIRT^71^. (5) fMRISurface for projecting the normalized functional data onto the standard CIFTI template. Additionally, the fMRI volumes are mapped from the cortical gray matter ribbon onto the native cortical surface and resampled to the 32k Conte69 surface template. These surfaces also are combined with the subcortical and cerebellar volumes into CIFTI format using the Connectome Workbench^72^. The surface and volumetric timeseries are smoothed with a 2 mm FWHM Gaussian kernel using geodesic and euclidean distances, respectively.

### DCAN BOLD Preprocessing

Standard fMRI BOLD preprocessing was further carried out using additional stages (Preproc) from the ABCD-BIDS Pipeline developed by the DCAN Labs^31,73^. The resting-state fMRI data were demeaned and detrended with respect to time. A general linear model was then created to denoise the processed fMRI data with regressors corresponding to signal and movement variables. The nuisance signal regressors comprised mean timeseries for the white matter, CSF, and mean CIFTI timeseries signal (proxy for the global signal). The white matter and CSF segmentations were derived during PostFreeSurfer. The nuisance motion regressors comprised translational (***x***, ***y***, *z*) and rotational (roll, pitch, yaw) parameters estimated by realignment during fMRIVolume and their Volterra expansion. After denoising the fMRI data, a bandpass filtering (0.008 < ***f*** < 0.09 Hz) was applied to the timeseries using a 2^nd^ order Butterworth filter. Physiological data also were acquired to derive the respiratory signal of each participant. A respiratory motion filter was applied to the motion realignment data to filter frequencies (18.582 to 25.726 breaths per minute) from the respiratory signal, improving the estimates of framewise displacement (FD)^18^ that were calculated during fMRIVolume. As mentioned in **Participants**, all the youth who were included in this study satisfied the structural T1-weighted and functional MRI data quality control criteria.

### Head Motion

Instantaneous frame displacement (FD) for a given frame *i* can be expressed as the sum of frame-to-frame change over the 6 rigid body (3 translational and 3 rotational) parameters^18,74^. Hence,

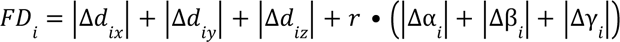

where Δ*d*_*ix*_ = *d*_(*i*−1)*x*_ − *d*_*ix*_, and similarly for the remaining rigid body parameters. We assigned *r* = 50 mm which approximates the mean distance from the cerebral cortex to the center of the head in neurotypical youth and adults^18,74^. Since FD_*i*_ is calculated by backward difference, it is computed for all the frames in an fMRI scan except the first one which is assigned a value of 0. For each participant, a mean FD can then be generated per fMRI scan by averaging their FD values. To identify “low-motion” youth in the ABCD Study, we applied a head motion threshold of mean FD < 0.20 mm to the participants’ resting-state fMRI data across a maximum of 4 runs. This threshold provides a reasonable balance between the sample size requirement to achieve adequate statistical power for brain-behavior associations and good data quality standards to mitigate motion artifacts in such studies. After applying a mean FD < 0.20 mm, a total of 3,342 youth remained in the ABCD Study, corresponding to ∼58% of the original sample size. This procedure disproportionately retained low-motion youth in the Black and Hispanic groups relative to the White group: White *N* = 2,266, Black *N* = 410, and Hispanic *N* = 666. The low-motion cohort comprised 61.6% White, 50.6% Black, and 53.4% Hispanic youth from their respective total sample sizes. Of note, a stricter mean FD threshold (e.g., mean FD > 0.08 mm) leads to greater data quality but at the expense of a drastic drop in sample size^1^.

### Parcellation Scheme

We used a whole-brain parcellation scheme to extract the mean timeseries of the youth across the 3 racial/ethnic groups for their resting-state fMRI scans^31,32^. This parcellation scheme extends the Gordon atlas that describes 333 cortical ROIs^32^ to include the subcortical and cerebellar structures with an additional 19 FreeSurfer ROIs. These 19 ROIs correspond to the bilateral nucleus accumbens, amygdala, caudate, hippocampus, pallidum, putamen, thalamus, ventral diencephalon, cerebellum, and extending to the midline brainstem. Although the properties of an individual’s functional connectome are better captured with cortical parcellation schemes at higher resolutions^2^, they are highly computationally intensive when constructing functional connectivity matrices in consortia datasets such as the ABCD Study. Further, whole-brain parcellations preserve the brain-behavior relationships for the NIH Toolbox and CBCL that are distributed across the subcortex and cerebellum^1,75^.

### Functional Connectivity with Standard Method

After extracting and concatenating the mean timeseries of each ROI from the extended Gordon parcellation across a maximum of 4 resting-state fMRI scans, the functional connectivity matrices of the youth were computed. For each participant, the Pearson correlation coefficient (*r*) was computed between all possible ROI pairs and Fisher *z*-transformed to construct a 352 x 352 functional connectivity matrix. The correlation values represent the connectivity strengths (edges) between two ROIs (nodes). For each youth, we removed duplicate edges across the whole brain by considering only the upper triangular part of their functional connectivity matrices. When the extended Gordon parcellation was applied, a total of [(352 x 352) – 352 / 2] = 61,776 unique edges were obtained in the functional connectome.

### Functional Connectivity with Scrubbing

To identify motion-corrupted timepoints in the fMRI timeseries of each youth, we applied a frame censoring strategy commonly known as scrubbing^14,18,33,34^ (see **Fig.1** for a schematic). For each youth, a motion-corrupted timepoint *T* is identified in their fMRI timeseries if its FD > 0.20 mm. Once *T* has been identified, the preceding timepoint (*T* – 1) and two succeeding timepoints (*T* + 1, *T* + 2) are censored in the fMRI timeseries. By removing a total of four timepoints for each *T* in the fMRI timeseries, the residual motion and micro head movements that persist in the neighboring timepoints of *T* are minimized. We did not impose a criterion to retain the youth who had at least a predefined number of scrubbed timepoints given their long fMRI timeseries that were concatenated across a maximum of 4 resting-state scans. There is no clear consensus on the recommended proportion of scrubbed timepoints to be used in fMRI-related studies although it has been suggested to retain participants who have at least 25-50% of their full timeseries or a scan length > 5 min^1,14,18,74^. Of note, other variants of frame censoring have been proposed such as excluding timepoints below an FD threshold that are less than five contiguous frames between sequential censored frames^31^ or filtering timepoints that are ordered by increasing FD values to create low-motion sliding windows of timeseries data^74^.

### Functional Connectivity with Motion-ordered and Bagging Methods

We employed the motion-ordered and bagging methods as described in our previous work^29^. For the <u>motion-ordered</u> method, we selected a subset of the least motion-corrupted timepoints denoted by a predefined threshold referred to as minTP (see **Fig.1** for a schematic). Applying the minTP threshold is important to prevent youth from having varying number of timepoints in their scrubbed timeseries given that some participants exhibit greater head motion compared to others. We ranked the scrubbed timeseries by their least FD values (0 → 0.20 mm) and selected the top timepoints based on the minTP threshold. The functional connectivity between all pairs of ROIs were computed from the minTP-matched timepoints using Pearson correlation coefficient (*r*) and Fisher *z*-transformed for each participant. This generated a motion-ordered functional connectivity matrix for each participant. Of note, some participants may have more minimally corrupted timepoints (FD → 0) whereas other participants may have more varying number of least corrupted timepoints (0 < FD < 0.20). The motion-ordered approach would generate a subset of the lowest-motion timepoints that could be favorable for participants whose timepoints are minimally corrupted by motion when constructing functional connectivity matrices. Instead of generating a single subset of the lowest-motion timepoints for a participant, we can resample their least motion-corrupted timepoints *with* replacement by creating multiple subsets of resampled timeseries data over a number of iterations to construct functional connectivity matrices − a procedure known as bagging^29^. Although the motion-ordered and bagging methods produced brain-behavior relationships that were equivocal in our previous proof-of-concept study^29^, which were somewhat limited in terms of the sample size, it remains crucial to fully assess whether bagging would have benefits in a larger, more diverse sample with more challenging levels of motion.

Next, we performed <u>bagging</u> at the timeseries level^29^. Bagging has been applied previously to resample fMRI data *within* and *between* participants with short acquisition scans^76–80^. Participant-level resampling of training data and aggregation of model parameters/features have been shown to improve the accuracy and generalizability of brain-behavior predictive models for a range of cognitive domains^77,78^. However, in these studies, all the participant fMRI data were sampled to generate subsamples of training data. Here, instead, we selected a subset of the least motion-corrupted timepoints based on the minTP threshold (see **Fig.1** for a schematic)^29^. Subsequently, we generated 500 bootstrapped samples of size TP *with* replacement from the minTP-matched timepoints for each participant. We chose to produce 500 bootstraps instead of 1,000^1^ as they provide a good balance between capturing the sampling variability across the 3 racial/ethnic groups and generating these samples from consortia datasets such as the ABCD Study using reasonable computational resources^81^. At each iteration, the functional connectivity between all pairs of ROIs were computed using Pearson correlation coefficient (*r*) and Fisher *z*-transformed for each youth. This generated a bagged functional connectivity matrix for every participant at each iteration.

### Brain-behavior Relationships with Standard, Motion-ordered, and Bagging Methods

To detect the <u>standard</u> brain-behavior relationships across the low-motion youth, we computed the partial Spearman’s Rank correlation (*R*_s_) between functional connectivity and NIH Toolbox as well as CBCL externalizing and internalizing at the edge level using the full timeseries while adjusting for sex assigned at birth and head motion (indexed by mean FD). The edge that shared the strongest brain-behavior relationship (highest *R*_s_) was selected after applying the Benjamini-Hochberg False Discovery Rate (q = 0.05)^36^ to correct for multiple comparisons across 61,776 edges. The strongest *R*_s_ was chosen based on its magnitude rather than sign to prevent penalizing negatively correlated edges (e.g., an edge’s *R*_s_ = −0.16 is stronger than another edge’s *R*_s_ = 0.15). The step of selecting the strongest edge and its ROI-ROI pair helps to reduce the high computational costs of computing brain-behavior relationships with the ABCD Study. We then established the reproducibility of the standard brain-behavior relationships across the 3 racial/ethnic groups across different sample sizes^1,29^. Each sample size was bootstrapped *without* replacement over 500 iterations, where we randomly selected samples of the low-motion youth at 11 logarithmically-spaced intervals: White *N* ∈ {25, 39, 62, 97, 152, 238, 374, 586, 920, 1444, 2266}; Black *N* ∈ {25, 33, 44, 58, 77, 101, 134, 177, 234, 310, 410}; Hispanic *N* ∈ {25, 35, 48, 67, 93, 129, 179, 249, 345, 480, 666}. For each low-motion racial/ethnic group and behavior, the standard method computed 11 intervals x 500 bootstrap *N* samples = 5,500 correlations. For each sample size, the mean brain-behavior *R*_s_ and 95% CIs were plotted as a function of *N*.

To detect the <u>motion-ordered</u> and <u>bagged</u> brain-behavior associations across the low-motion youth, we repeated the analyses by computing *R*_s_ between functional connectivity and NIH Toolbox as well as CBCL externalizing and internalizing using the scrubbed timeseries while treating sex assigned at birth and head motion (indexed by mean FD) as covariates^29^. The same edge that shared the highest correlation strength with the standard method after applying the Benjamini-Hochberg False Discovery Rate (q = 0.05)^36^ was chosen for the motion-ordered and bagging procedures. The reproducibility of the brain-behavior *R*_s_ was computed based on the motion-ordered and bagged functional connectivity matrices across different sample sizes following similar sampling intervals as the standard method. For each low-motion racial/ethnic group and behavior, the motion-ordered method computed 11 intervals x 500 bootstrap *N* samples = 5,500 correlations. For each low-motion racial/ethnic group and behavior, the bagging procedure computed 11 intervals x 500 bootstrap *N* samples x 500 bootstrap timepoints = 2.75 million correlations. The mean brain-behavior *R*_s_ and 95% CIs were plotted as a function of *N*. We then assessed the performance of the standard, motion-ordered, and bagging methods by estimating their areas under the curve (AUC). To compare the difference in AUC between the standard and motion-ordered/bagging methods, we computed the rate of change in AUC as denoted by ΔAUC. In theory, an ΔAUC = 0 denotes identical effect sizes produced by the standard and motion-ordered or bagging methods. A positive ΔAUC reflects tighter 95% CIs produced by the motion-ordered or bagging method relative to the standard method whereas a negative ΔAUC corresponds to larger 95% CIs produced by the motion-ordered or bagging method relative to the standard method. However, the aim is to produce comparable or stronger brain-behavior effect sizes, where |ΔAUCs| are within an acceptance range. Therefore, AUC provides a simple, interpretable, and low computationally intensive metric to compare the effect sizes produced by the standard, motion-ordered, and bagging methods.

### Retaining High-Motion Youth for Inclusive Brain-Behavior Relationships

To increase participant inclusion in the ABCD Study, we examined the possibility of computing the brain-behavior *R*_s_ using the motion-ordered and bagged functional connectivity matrices derived from the scrubbed fMRI timeseries of youth *without* applying the initial mean FD < 0.20 mm^29^. That is, we retained all the youth across the 3 racial/ethnic groups who had scrubbed timepoints that matched the minTP threshold — including the “high-motion” participants who had a mean FD > 0.20 mm and who would have been discarded. The reproducibility of the brain-behavior *R*_s_ was recomputed on the basis of the motion-ordered and bagged functional connectivity matrices for different sample sizes derived from all the participants who had enough “usable” scrubbed timepoints across the 3 racial/ethnic groups. The mean brain-behavior *R*_s_ and 95% CIs also were plotted as a function of *N*. Finally, the performances of the motion-ordered and bagging methods were reassessed (using the AUC) and compared with the standard method (using the ΔAUC) when the eligible “high-motion” youth were included in the brain-behavior analyses. As described above, the aim is to produce comparable or stronger brain-behavior effect sizes, where the |ΔAUCs| are within an acceptance range as they are estimated by including the “high-motion” youth who typically are not retained in such studies.

### Concordance of Brain-Behavior Relationships from Low-/High-Motion Youth

We examined the concordance of the brain-behavior relationships obtained from the standard and motion-ordered/bagging methods. We computed the Pearson correlation coefficient (*r*) and Lin’s concordance correlation coefficient (ρ_c_)^37,38^ between the standard and motion-ordered/bagged brain-behavior relationships from the low-motion youth at typical (*N* = 25) and large (*N* = 250) sample sizes. We then repeated the analyses by including all the “high-motion” youth who were retained based on their least motion-corrupted timepoints that matched the minTP threshold. ρ_c_ quantifies the agreement between two measures obtained from a set of bivariate data ranging from −1 to 1. While it can be interpreted as a weighted version of Pearson’s *r*, two strongly correlated measures can still exhibit poor reproducibility if the mean squared error is high^38^. Hence, ρ_c_ will be high for two strongly correlated measures that have low mean squared error. If two measures are maximally correlated (*r* = 1), then ρ_c_ will be 1, otherwise, it will approximate to 0 (*r* → 0) or if the means or variances of the two population samples largely differ. However, ρ_c_ will be negative for two anticorrelated measures (*r* < 0) that have similar means and variances. ρ_c_ can be calculated for two samples *x*_1_and *x*_2_:

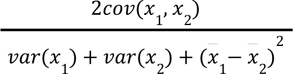

We also repeated the brain-behavior analyses derived from the first resting-state fMRI scan (REST1) instead of using a maximum of four scans given that community-based samples and consortia datasets do not always acquire multiple scans for each individual^39^. We aimed to reproduce the brain-behavior associations to assess whether the “high-motion” youth in the ABCD Study can still be retained on the basis of their motion-limited timeseries using typical scan duration (1 run x 5 min).

As all the brain-behavior associations were derived based on the edge that shared the *strongest R*_s_ in the whole-brain functional connectome, we reassessed the ability of the motion-ordered and bagging methods to reproduce the associations from the *weakest* edge that survived the Benjamini-Hochberg False Discovery Rate (q = 0.05)^36^ for each behavior. Finally, we selected the edge that exhibited the lowest *R*_s_ in terms of magnitude rather than sign to prevent penalizing connections that are negatively correlated with the behaviors. An edge may exhibit high (low) test-retest reliability but share low (high) behavioral relevance. This is because the most (least) reliable edge in the functional connectome may not (may) contain useful information for predicting or relating to behavior^40^.

## Data Availability

The ABCD Study is openly available following access permission granted to one or multiple NIMH Data Archive (NDA) Collections (https://nda.nih.gov/nda/access-data-info). The ABCD data repository grows and changes over time (https://nda.nih.gov/). The ABCD data used in this report came from the ABCD Collection 3165 and preprocessed using the ABCD-BIDS Community Collection from the DCAN Labs (https://nda.nih.gov/edit_collection.html?id=3165).

## Code Availability

The analysis code for the motion-ordered and bagging methods can be found at https://github.com/JRam02/bagging. The analysis code specific to this study can be found at https://github.com/JRam02/inclusivity. The code for processing the ABCD Study by the ABCD-BIDS Community Collection can be found at https://github.com/DCAN-Labs/abcd-hcp-pipeline. The MRI data analysis code can be found at https://github.com/ABCD-STUDY/nda-abcd-collection-3165.

## Acknowledgments

Data used in the preparation of this article were obtained from the ABCD Study^®^ (<u>abcdstudy.org/</u>), held in the NIMH Data Archive (NDA). This is a multisite, longitudinal study designed to recruit more than 10,000 children aged 9-10 and follow them over 10 years into early adulthood. The ABCD Study is supported by the National Institutes of Health (NIH) and additional federal partners under award numbers: U01DA041048, U01DA050989, U01DA051016, U01DA041022, U01DA051018, U01DA051037, U01DA050987, U01DA041174, U01DA041106, U01DA041117, U01DA041028, U01DA041134, U01DA050988, U01DA051039, U01DA041156, U01DA041025, U01DA041120, U01DA051038, U01DA041148, U01DA041093, U01DA041089, U24DA041123, U24DA041147. The full list of federal supporters is available at https://abcdstudy.org/federal-partners.html. The complete lists of participating sites and study investigators can be found at https://abcdstudy.org/consortium_members/. The ABCD Consortium investigators designed and implemented the study and/or provided the data but did not necessarily participate in the analysis or writing of this report. This manuscript reflects the views of the authors and may not reflect the opinions or views of the NIH or ABCD Consortium investigators. Additional support for this work was made possible from NIEHS R01-ES032295, R01-ES031074, and R21DA057592. This work also obtained support from the Yale Kavli Institute for Neuroscience and the Wu Tsai Institute at Yale University. This work used the computational resources from the Masonic Institute for the Developing Brain (MIDB), Neuroimaging Genomics Data Resource (NGDR), and Minnesota Supercomputing Institute (MSI) at the University of Minnesota. We thank the ABCD JEDI Workgroup 3 (Responsible Use of ABCD Study Data) for the discussions regarding the motivations and findings of this study.

## Author Contributions Statement

J.R developed, tested, and produced the brain-behavior analyses. J.R and C.K developed the motion-ordered and bagging methods. J.R and A.B.-S conceptualized this study, interpreted the analyses, and wrote the original draft of the manuscript. L.Q.U, T.V, and C.K assisted in interpreting the analyses as well as writing and reviewing the manuscript. E.F and D.A.F provided access to the ABCD-BIDS Community Collection data and computational resources from the Masonic Institute for the Developing Brain (MIDB), Neuroimaging Genomics Data Resource (NGDR), and Minnesota Supercomputing Institute (MSI) at the University of Minnesota. A.B.-S supervised this study.

## Competing Interests Statement

D.A.F has a financial interest in Turing Medical Inc. and may benefit financially if the company is successful in marketing FIRMM motion monitoring software products. D.A.F may receive royalty income based on FIRMM technology developed at Washington University School of Medicine and Oregon Health and Sciences University and licensed to Turing Medical Inc. D.A.F is a co-founder of Turning Medical Inc. These potential conflicts of interest have been reviewed and are managed by Washington University School of Medicine, Oregon Health and Sciences University, and the University of Minnesota. The other authors declare no competing interests.

