## Supplementary Information for "Increasing the representation of minoritized youth for inclusive and reproducible brain-behavior associations"

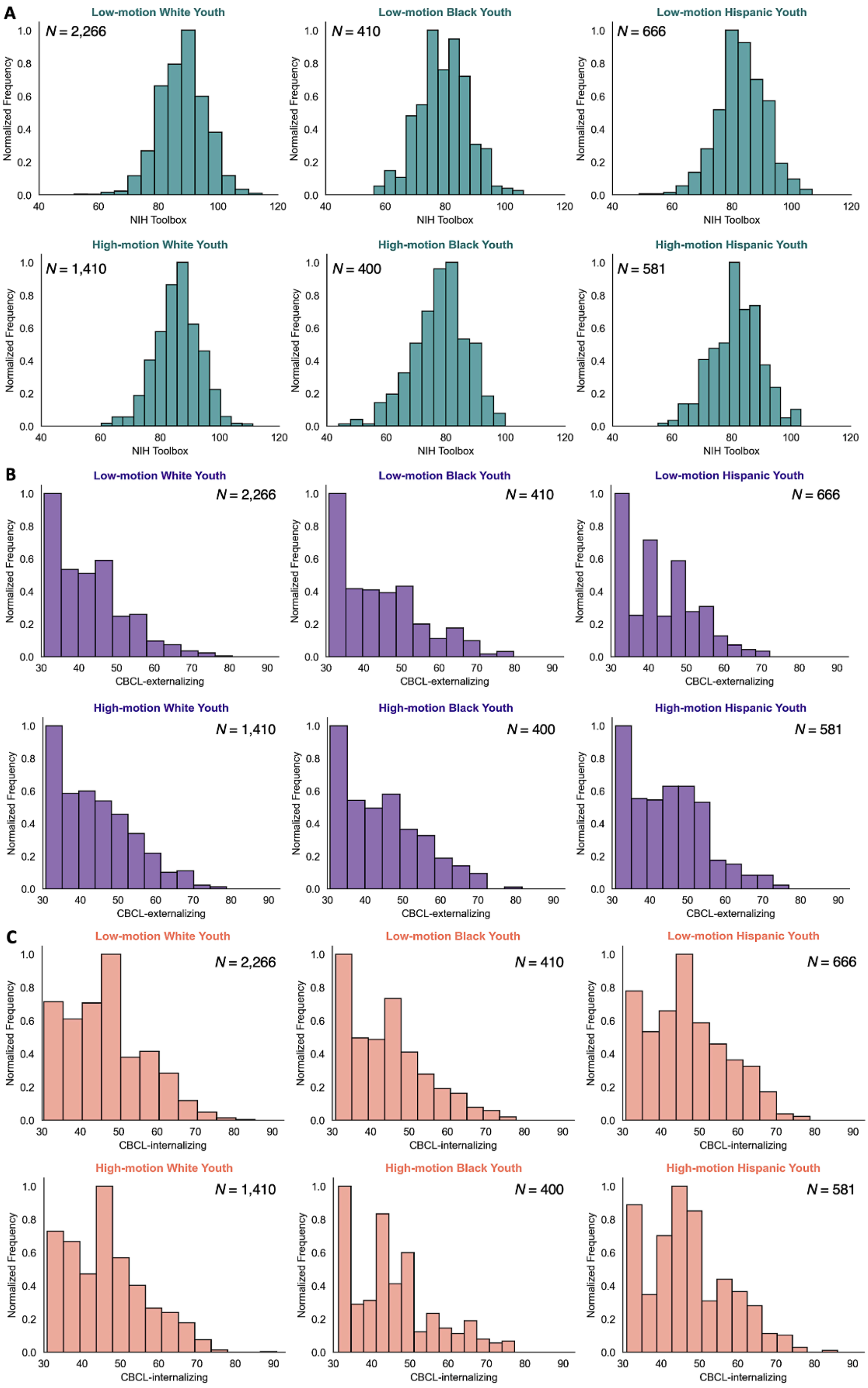


**Supplementary Figure 1. Distributions of the (A) NIH Toolbox, (B) CBCL-externalizing, and (C) CBCL-internalizing across the 3 racial/ethnic groups.** There was a significance in the NIH Toolbox distributions between the low-motion and high-motion White and Hispanic youth (two-sample Kolmogorov Smirnov test, White D = 0.13, $P < 0.001$; Hispanic D = 0.12, $P < 0.001$). There was a significance in the CBCL-externalizing distributions between the low-motion and high-motion White youth (two-sample Kolmogorov Smirnov test, D = 0.075, $P < 0.001$). There were no significant differences in the CBCL-internalizing distributions between the low-motion and high-motion racial/ethnic groups (two-sample Kolmogorov Smirnov test, $P > 0.05$).

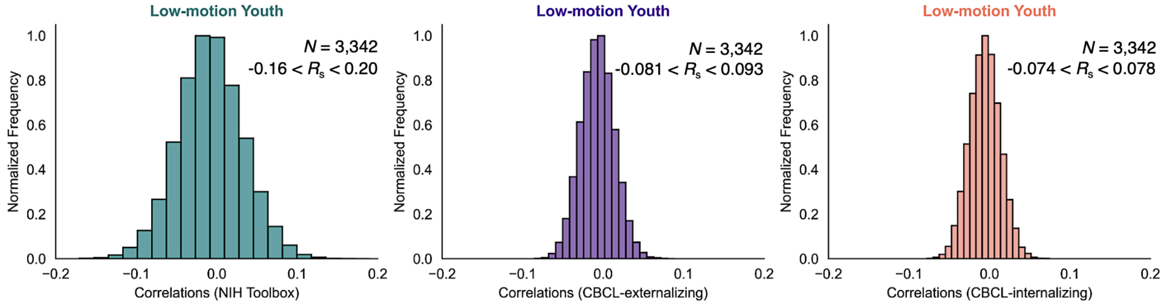


**Supplementary Figure 2. Distributions of the edge-level correlations of the NIH Toolbox, CBCL-externalizing, and CBCL-internalizing derived from the low-motion youth.** Partial Spearman's Rank correlations ($R_s$) were computed between functional connectivity and the three behaviors at the edge-level while treating sex assigned at birth and mean FD as covariates derived from the low-motion youth across the 3 racial/ethnic groups. This allowed us to obtain the edge that shared the strongest brain-behavior relationship ($R_s$) for each behavior that was consistent across the 3 low-motion racial/ethnic groups after applying the Benjamini-Hochberg False Discovery Rate (q = 0.05) to correct for multiple comparisons across 61,776 edges. For each behavior, the range of $R_s$ is displayed across 61,776 edges in the functional connectome before correcting for multiple comparisons. Of note, 17,854 and 65 respective edges survived multiple comparison corrections for the NIH Toolbox and CBCL-externalizing. However, there was no edge in the functional connectome that survived statistical significance after correcting for multiple comparisons for CBCL-internalizing.

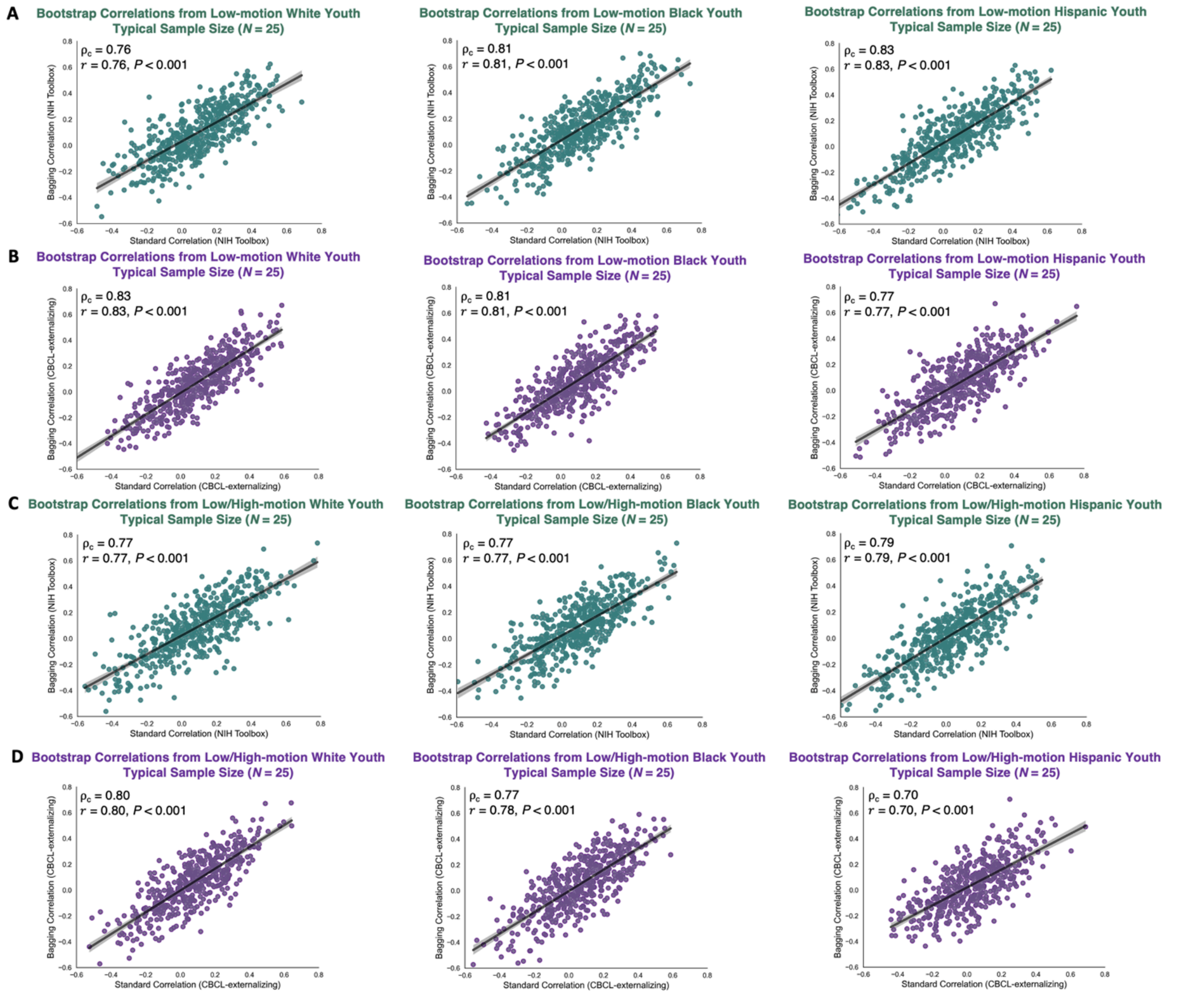


**Supplementary Figure 3. Bootstrap brain-behavior correlations obtained from the standard and bagged methods at typical sample size ($N$ = 25) *with* (A-B) and *without* (C-D) excluding the high-motion youth across the 3 racial/ethnic groups.** The teal and purple data points corresponded to bootstrap samples at $N$ = 25 obtained from 500 iterations and these samples were used to compute the associations between functional connectivity and NIH Toolbox and CBCL-externalizing. The Lin's concordance correlation coefficient ($\rho_c$) and Pearson correlation coefficient ($r$) were computed respectively to assess the reproducibility and similarity of the brain-behavior associations obtained from the standard and bagging methods. The low-/high-motion youth were retained based on the assumption that they had a minimum of 100 least motion-corrupted timepoints in their fMRI timeseries.

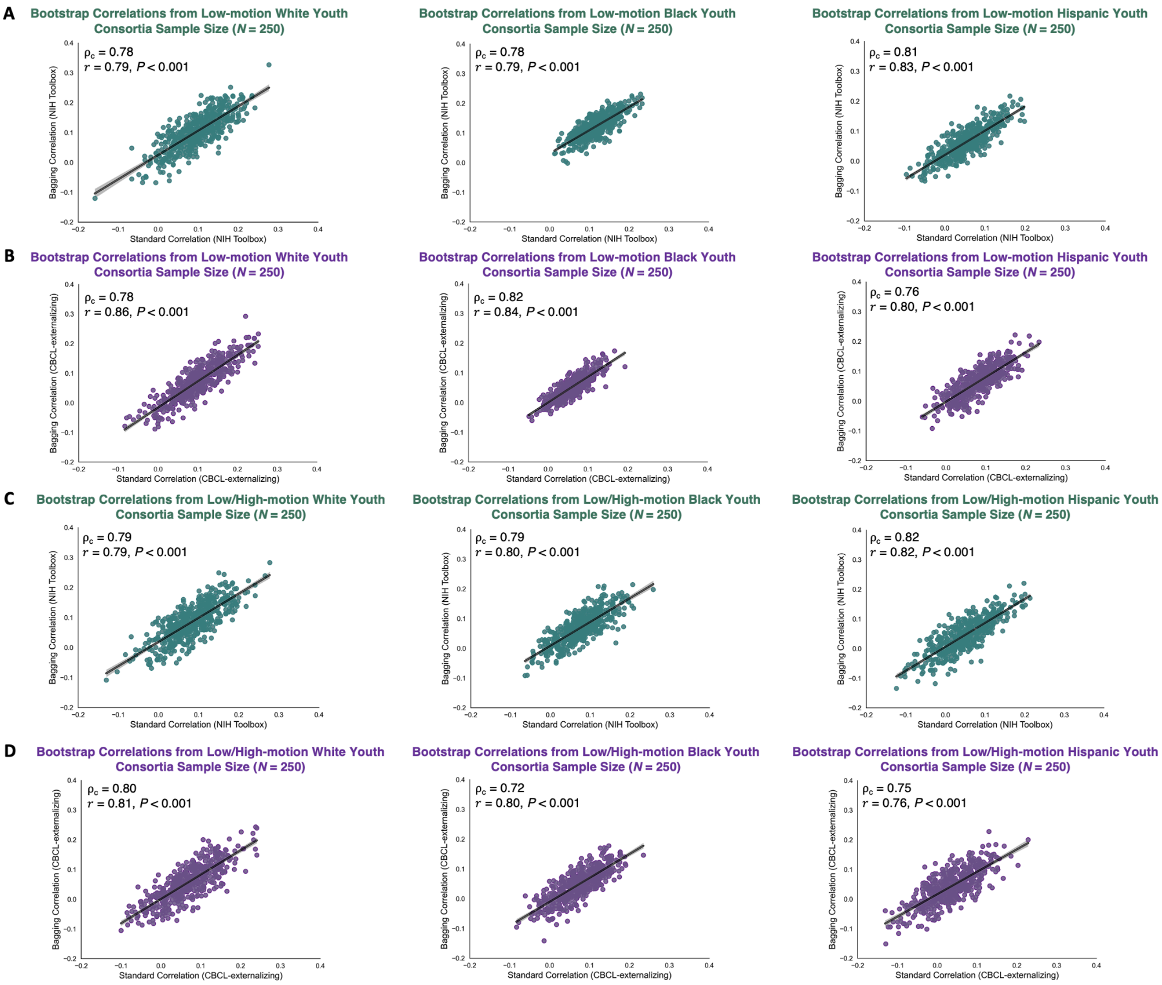


**Supplementary Figure 4. Bootstrap brain-behavior correlations obtained from the standard and bagging methods at large sample size ($N$ = 250) *with* (A-B) and *without* (C-D) excluding the high-motion youth across the 3 racial/ethnic groups.** The teal and purple data points corresponded to bootstrap samples at $N$ = 250 obtained from 500 iterations and these samples were used to compute the associations between functional connectivity and NIH Toolbox and CBCL-externalizing. The Lin's concordance correlation coefficient ($\rho_c$) and Pearson correlation coefficient ($r$) were computed respectively to assess the reproducibility and similarity of the brain-behavior associations obtained from the standard and bagging methods. The low-/high-motion youth were retained based on the assumption that they had a minimum of 100 least motion-corrupted timepoints in their fMRI timeseries.

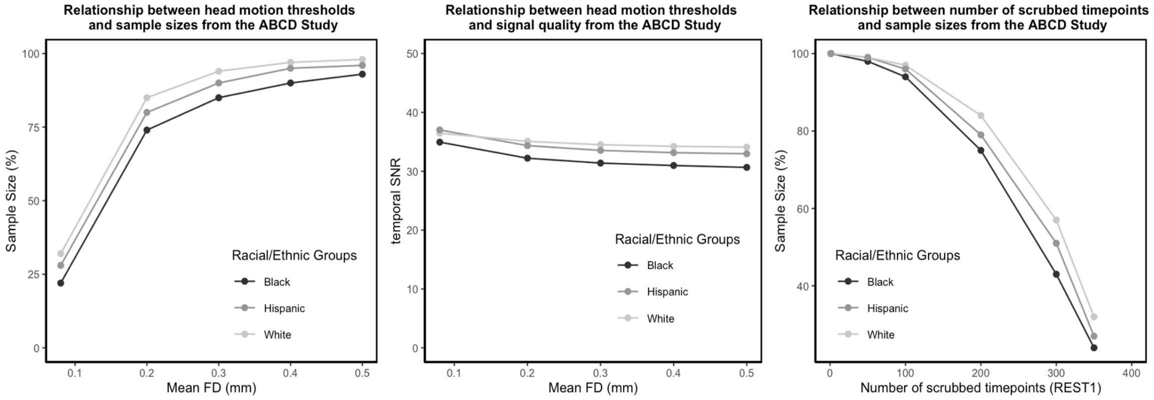


**Supplementary Figure 5. Impact of head motion derived from REST1 across racial and ethnic groups in the ABCD Study.** Left: Relationship between head motion thresholds and sample sizes. Head motion was quantified by mean framewise displacement (FD) such that mean FD $\in \{0.08, 0.20, 0.30, 0.40, 0.50\}$. Middle: Relationship between head motion thresholds and signal quality indexed by the temporal signal-to-noise ratio (tSNR). Mean FD $\in \{0.08, 0.20, 0.30, 0.40, 0.50\}$. Right: Relationship between number of scrubbed timepoints and sample sizes. Number of scrubbed timepoints $\in \{1, 50, 100, 200, 300, 350\}$.

| ABCD Study | Racial/Ethnic Groups | | |
|---|---|---|---|
| | **White** | **Black** | **Hispanic** |
| Scans | Baseline | Baseline | Baseline |
| Sample Size | 3,676 | 810 | 1,247 |
| Age (years) | 9-10 | 9-10 | 9-10 |
| ADI[a] | 1.1-125.7 | 1.1-124.5 | 1.1-124.6 |
| Sex[b] (Female:Male) (%) | 48:52 | 53:47 | 50:50 |
| NIH Toolbox[c] | 54-117 | 46-108 | 51-109 |
| CBCL-Externalizing[d] | 33-83 | 33-84 | 33-79 |
| CBCL-Internalizing[d] | 33-93 | 33-80 | 33-88 |
| Sites | 21 | 21 | 21 |
| Frames[e] | 380-1,570 | 196-1,560 | 542-1,560 |
| tSNR[f] | 10.8-55.6 | 11.7-49.8 | 11.2-53.0 |
| Mean FD[g] (mm) | 0.03-2.4 | 0.04-2.1 | 0.04-2.2 |

**Supplementary Table 1. Demographic, behavioral, and fMRI characteristics derived from the ABCD Study NIMH Data Archive Release 4.0.** [a]ADI = Area Deprivation Index. [b]Participant sex denoted youth's sex assigned at birth. [c]NIH Toolbox = National Institutes of Health Toolbox indexing cognitive performance. [d]CBCL = Child Behavior Checklist indexing externalizing and internalizing behaviors. [e]Frames = number of volumes acquired across 4 resting-state scans. [f]tSNR = temporal signal-to-noise ratio. [g]FD = Framewise Displacement. Note that the externalizing and internalizing CBCL and NIH Toolbox scores were $T$-standardized.

The effect sizes produced by the standard and motion-ordered methods from REST1 were comparable for the NIH Toolbox (0.20% < ΔAUC < 5.03%) and CBCL-externalizing (1.31% < ΔAUC < 4.06%) across the 3 low-motion racial/ethnic groups.

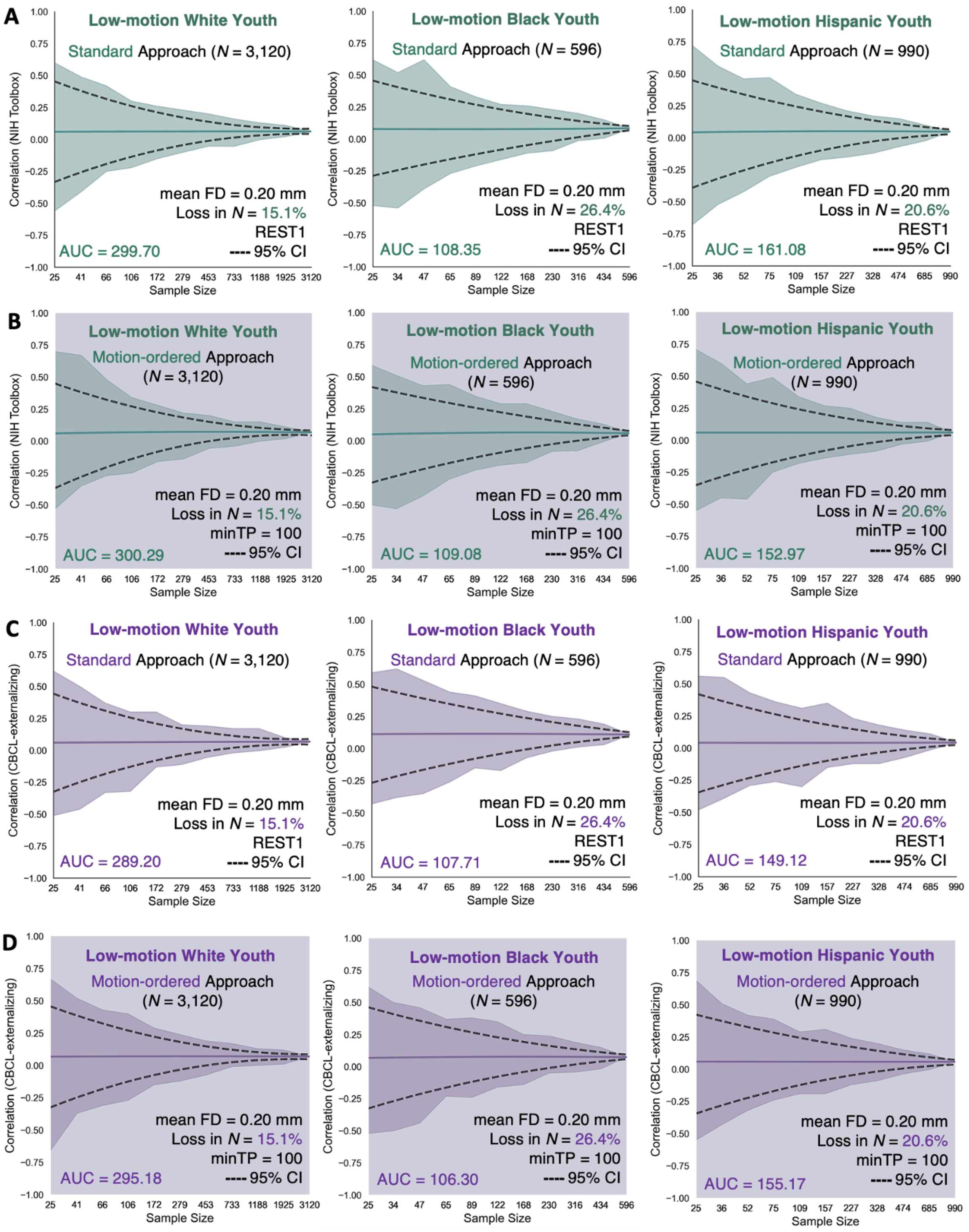


**Supplementary Figure 6. Correlations between functional connectivity and (A-B) NIH Toolbox and (C-D) CBCL-externalizing using the standard and motion-ordered methods as a function of sample size across the 3 low-motion racial/ethnic groups obtained from REST1.** The respective edge that was chosen in the functional connectome derived from a single fMRI run (REST1) shared the strongest correlation strength with the NIH Toolbox and CBCL-externalizing. The standard method corresponded to the brain-behavior associations derived from the full fMRI timeseries of the low-motion youth with a mean FD < 0.20 mm. The motion-ordered method corresponded to the brain-behavior associations derived from the scrubbed fMRI timeseries ranked and thresholded by their 100 least motion-corrupted timepoints ($0 < FD < 0.20$ mm) to construct the functional connectivity matrices of the low-motion youth. The sample sizes were bootstrapped at 11 logarithmically-spaced $N$ intervals: White $N \in \{25, 41, 66, 106, 172, 279, 453, 733, 1188, 1925, 3120\}$; Black $N \in \{25, 34, 47, 65, 89, 122, 168, 230, 316, 434, 596\}$; Hispanic $N \in \{25, 36, 52, 75, 109, 157, 227, 328, 474, 685, 990\}$. Solid teal and purple lines show the mean correlations from the 500 bootstrap samples for a given same size. Teal and purple shadings denote the minimum and maximum correlations across 500 bootstrap samples for a given sample size. Black dotted lines represent the lower and upper bounds of the 95% CIs for a given sample size. The areas under the curve (AUC) for the brain-behavior associations also are displayed.

The effect sizes produced by the standard and motion-ordered methods from REST1 remained comparable for the NIH Toolbox (1.00% < ΔAUC < 3.91%) and CBCL-externalizing (1.34% < ΔAUC < 2.18%) across the 3 racial/ethnic groups when the high-motion youth were included.

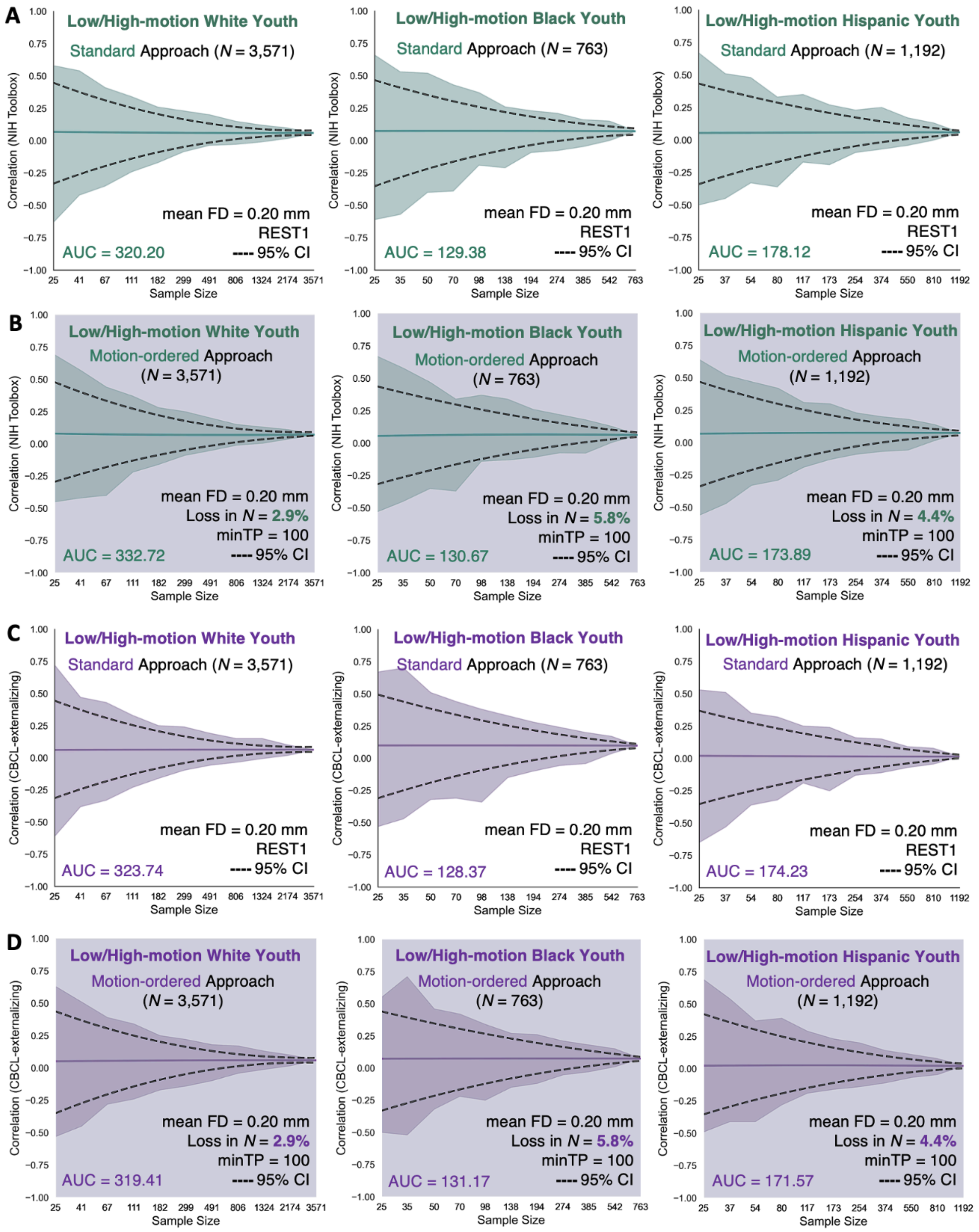


**Supplementary Figure 7. Correlations between functional connectivity and (A-B) NIH Toolbox and (C-D) CBCL-externalizing using the standard and motion-ordered methods as a function of sample size across the 3 low-/high-motion racial/ethnic groups obtained from REST1.** The respective edge that was chosen in the functional connectome derived from a single fMRI run (REST1) shared the strongest correlation strength with the NIH Toolbox and CBCL-externalizing. The standard method corresponded to the brain-behavior associations derived from the full fMRI timeseries of the low-/high-motion youth that have been retained for the analyses without imposing an initial head motion threshold of mean FD < 0.20 mm. The motion-ordered method corresponded to the brain-behavior associations derived from the scrubbed fMRI timeseries of the low-/high-motion youth who were retained based on the assumption that they had a minimum of 100 least motion-corrupted timepoints. Their scrubbed fMRI timeseries were ranked by their lowest FD values and 100 least motion-corrupted timepoints (0 < FD < 0.20 mm) were selected to construct the functional connectivity matrices of the youth. The sample sizes were bootstrapped at 11 logarithmically-spaced *N* intervals: White $N \in$ {25, 41, 67, 111, 182, 299, 491, 806, 1324, 2174, 3571}; Black $N \in$ {25, 35, 50, 70, 98, 138, 194, 274, 385, 542, 763}; Hispanic $N \in$ {25, 37, 54, 80, 117, 173, 254, 374, 550, 810, 1192}. Solid teal and purple lines show the mean correlations from the 500 bootstrap samples for a given same size. Teal and purple shadings denote the minimum and maximum correlations across 500 bootstrap samples for a given sample size. Black dotted lines represent the lower and upper bounds of the 95% CIs for a given sample size. The areas under the curve (AUC) for the brain-behavior associations also are displayed.

The effect sizes produced by the standard and bagging methods from REST1 were comparable for the NIH Toolbox (0.30% < ΔAUC < 7.15%) and CBCL-externalizing (0.45% < ΔAUC < 3.78%) across the 3 low-motion racial/ethnic groups.

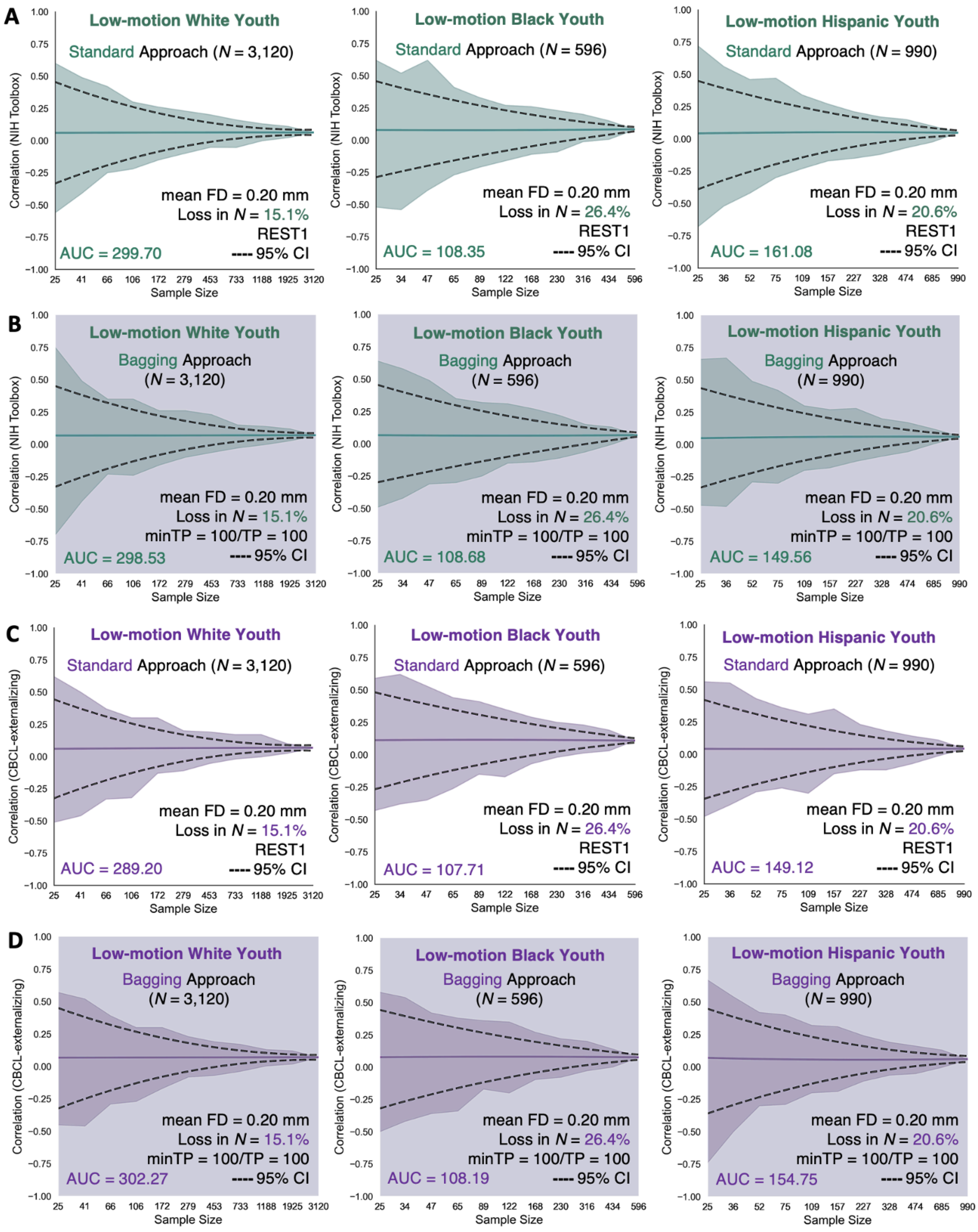


**Supplementary Figure 8. Correlations between functional connectivity and (A-B) NIH Toolbox and (C-D) CBCL-externalizing using the standard and bagging methods as a function of sample size across the 3 low-motion racial/ethnic groups obtained from REST1.** The respective edge that was chosen in the functional connectome derived from a single fMRI run (REST1) shared the strongest correlation strength with the NIH Toolbox and CBCL-externalizing. The standard method corresponded to the brain-behavior associations derived from the full fMRI timeseries of the low-motion youth with a mean FD < 0.20 mm. The bagging method corresponded to the brain-behavior associations derived from the scrubbed fMRI timeseries ranked and thresholded by their 100 least motion-corrupted timepoints ($0 < FD < 0.20$ mm) from which 100 timepoints were bootstrapped across 500 iterations to construct the functional connectivity matrices of the low-motion youth. The sample sizes were bootstrapped at 11 logarithmically-spaced *N* intervals: White $N \in \{25, 41, 66, 106, 172, 279, 453, 733, 1188, 1925, 3120\}$; Black $N \in \{25, 34, 47, 65, 89, 122, 168, 230, 316, 434, 596\}$; Hispanic $N \in \{25, 36, 52, 75, 109, 157, 227, 328, 474, 685, 990\}$. Solid teal and purple lines show the mean correlations from the 500 bootstrap samples for a given same size. Teal and purple shadings denote the minimum and maximum correlations across 500 bootstrap samples for a given sample size. Black dotted lines represent the lower and upper bounds of the 95% CIs for a given sample size. The areas under the curve (AUC) for the brain-behavior associations also are displayed.

The effect sizes produced by the standard and bagging methods from REST1 remained comparable for the NIH Toolbox (0.58% < ΔAUC < 4.90%) and CBCL-externalizing (0.37% < ΔAUC < 0.90%) across the 3 racial/ethnic groups when the high-motion youth were included.

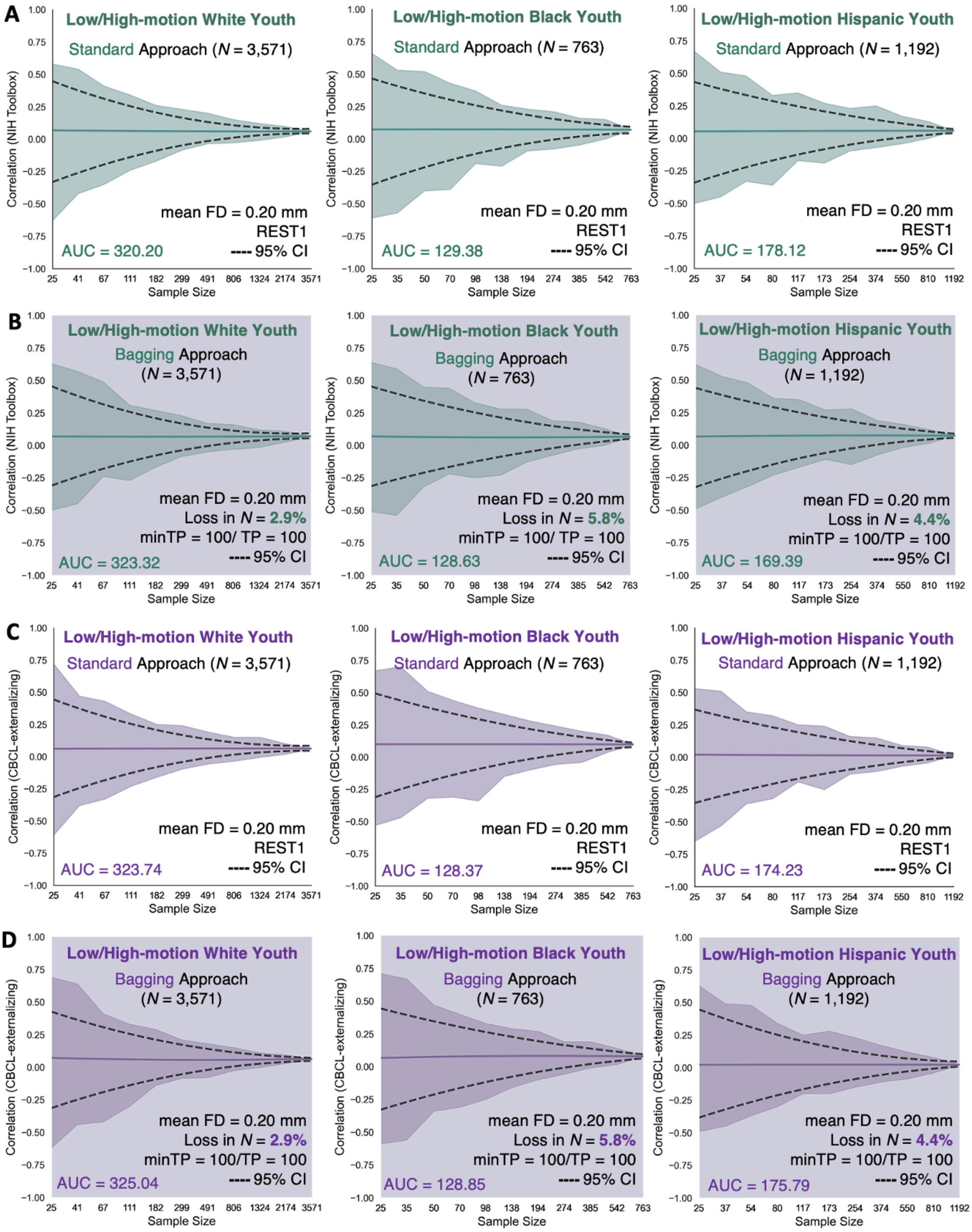


**Supplementary Figure 9. Correlations between functional connectivity and (A-B) NIH Toolbox and (C-D) CBCL-externalizing using the standard and bagging methods as a function of sample size across the 3 low-/high-motion racial/ethnic groups obtained from REST1.** The respective edge that was chosen in the functional connectome derived from a single fMRI run (REST1) shared the strongest correlation strength with the NIH Toolbox and CBCL-externalizing. The standard method corresponded to the brain-behavior associations derived from the full fMRI timeseries of the low-/high-motion youth that have been retained for the analyses without imposing an initial head motion threshold of mean FD < 0.20 mm. The bagging method corresponded to the brain-behavior associations derived from the scrubbed fMRI timeseries of the low-/high-motion youth who were retained based on the assumption that they had a minimum of 100 least motion-corrupted timepoints. Their scrubbed fMRI timeseries were ranked by their lowest FD values and 100 least motion-corrupted timepoints ($0 < FD < 0.20$ mm) were selected from which 100 timepoints were bootstrapped across 500 iterations to construct the functional connectivity matrices of the youth. The sample sizes were bootstrapped at 11 logarithmically-spaced *N* intervals: White $N \in \{25, 41, 67, 111, 182, 299, 491, 806, 1324, 2174, 3571\}$; Black $N \in \{25, 35, 50, 70, 98, 138, 194, 274, 385, 542, 763\}$; Hispanic $N \in \{25, 37, 54, 80, 117, 173, 254, 374, 550, 810, 1192\}$. Solid teal and purple lines show the mean correlations from the 500 bootstrap samples for a given same size. Teal and purple shadings denote the minimum and maximum correlations across 500 bootstrap samples for a given sample size. Black dotted lines represent the lower and upper bounds of the 95% CIs for a given sample size. The areas under the curve (AUC) for the brain-behavior associations also are displayed.

The effect sizes produced by the standard and motion-ordered methods using the weakest edge in the functional connectome were comparable for the NIH Toolbox (1.45% < ΔAUC < 5.19%) and CBCL-externalizing (0.44% < ΔAUC < 6.06%) across the 3 low-motion racial/ethnic groups.

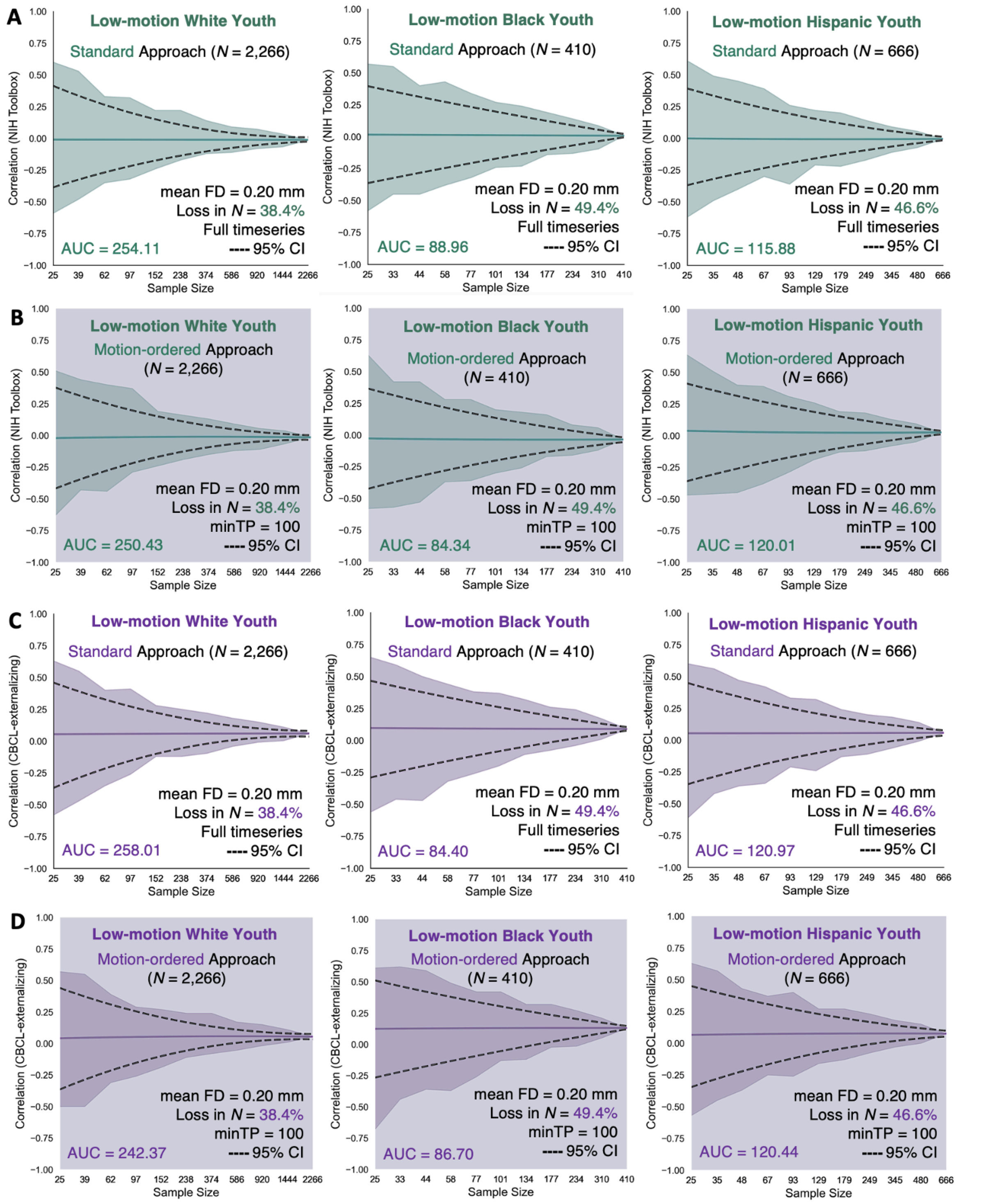


**Supplementary Figure 10. Correlations between functional connectivity and (A-B) NIH Toolbox and (C-D) CBCL-externalizing using the standard and motion-ordered methods as a function of sample size across the 3 low-motion racial/ethnic groups obtained from the weakest edge in the functional connectome.** The respective edge that was chosen in the functional connectome shared the weakest correlation strength with the NIH Toolbox ($|R_s|$ = 0.042, $P$ < 0.05 FDR) and CBCL-externalizing ($|R_s|$ = 0.070, $P$ < 0.05 FDR). The standard method corresponded to the brain-behavior associations derived from the full fMRI timeseries of the low-motion youth with a mean FD < 0.20 mm. The motion-ordered method corresponded to the brain-behavior associations derived from the scrubbed fMRI timeseries ranked and thresholded by their 100 least motion-corrupted timepoints ($0 < FD < 0.20$ mm) to construct the functional connectivity matrices of the low-motion youth. The sample sizes were bootstrapped at 11 logarithmically-spaced *N* intervals: White $N \in \{25, 39, 62, 97, 152, 238, 374, 586, 920, 1444, 2266\}$; Black $N \in \{25, 33, 44, 58, 77, 101, 134, 177, 234, 310, 410\}$; Hispanic $N \in \{25, 35, 48, 67, 93, 129, 179, 249, 345, 480, 666\}$. Solid teal and purple lines show the mean correlations from the 500 bootstrap samples for a given same size. Teal and purple shadings denote the minimum and maximum correlations across 500 bootstrap samples for a given sample size. Black dotted lines represent the lower and upper bounds of the 95% CIs for a given sample size. The areas under the curve (AUC) for the brain-behavior associations also are displayed.

The standard and motion-ordered effect sizes that were obtained from the weakest edge in the functional connectome remained comparable for the NIH Toolbox (0.25% < ΔAUC < 1.31%) and CBCL-externalizing (0.30% < ΔAUC < 6.66%) across the 3 racial/ethnic groups when the high-motion youth were included.

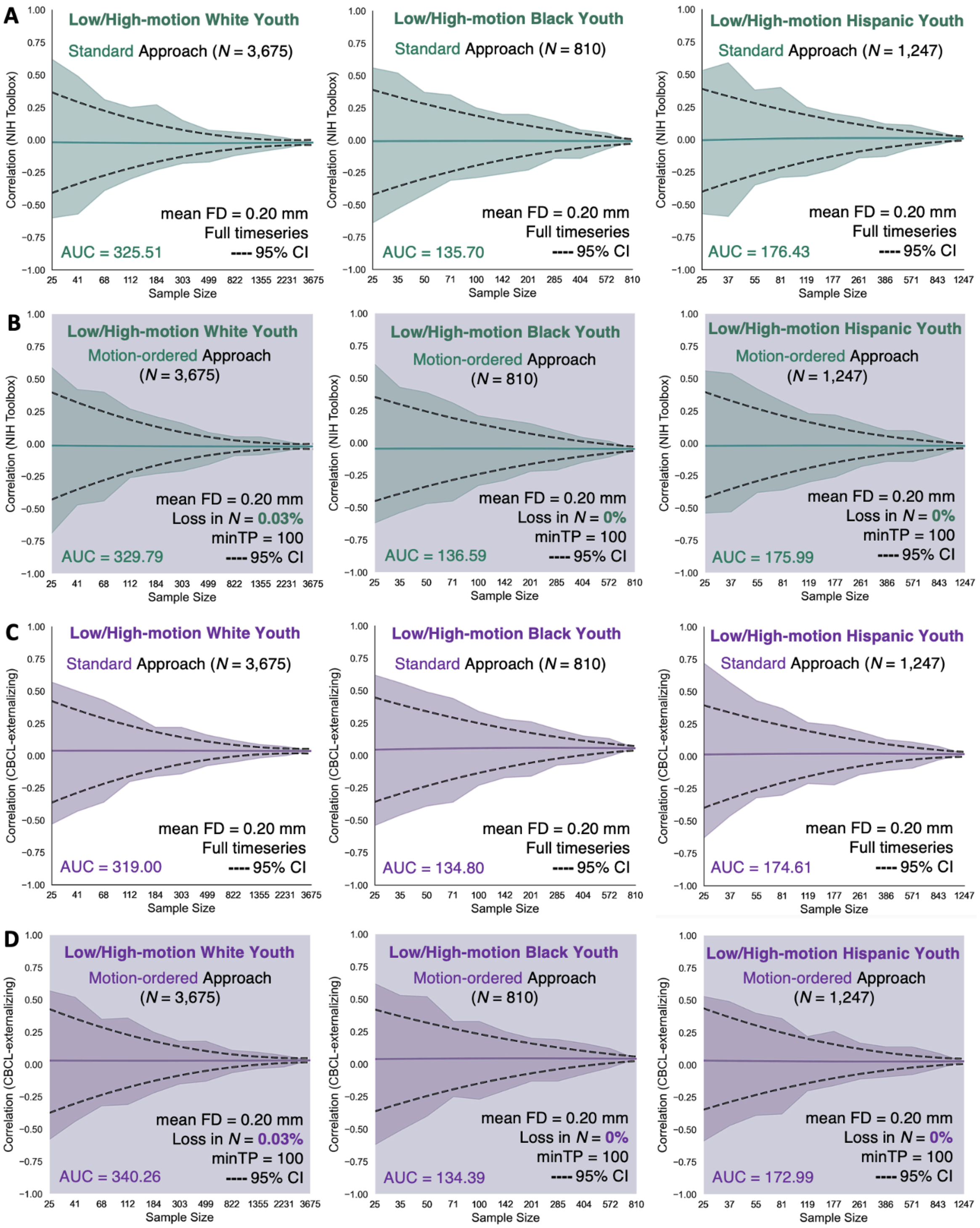


**Supplementary Figure 11. Correlations between functional connectivity and (A-B) NIH Toolbox and (C-D) CBCL-externalizing using the standard and motion-ordered methods as a function of sample size across the 3 low/high-motion racial/ethnic groups obtained from the weakest edge in the functional connectome.** The respective edge that was chosen in the functional connectome shared the weakest correlation strength with the NIH Toolbox ($|R_s|$ = 0.042, $P < 0.05$ FDR) and CBCL-externalizing ($|R_s|$ = 0.070, $P < 0.05$ FDR). The standard method corresponded to the brain-behavior associations derived from the full fMRI timeseries of the low-/high-motion youth that have been retained for the analyses without imposing an initial head motion threshold of mean FD < 0.20 mm. The motion-ordered method corresponded to the brain-behavior associations derived from the scrubbed fMRI timeseries of the low-/high-motion youth who were retained based on the assumption that they had a minimum of 100 least motion-corrupted timepoints. Their scrubbed fMRI timeseries were ranked by their lowest FD values and 100 least motion-corrupted timepoints (0 < FD < 0.20 mm) were selected to construct the functional connectivity matrices of the youth. The sample sizes were bootstrapped at 11 logarithmically-spaced *N* intervals: White $N \in \{25, 41, 68, 112, 184, 303, 499, 822, 1355, 2231, 3675\}$; Black $N \in \{25, 35, 50, 71, 100, 142, 201, 285, 404, 572, 810\}$; Hispanic $N \in \{25, 37, 55, 81, 119, 177, 261, 386, 571, 843, 1247\}$. Solid teal and purple lines show the mean correlations from the 500 bootstrap samples for a given same size. Teal and purple shadings denote the minimum and maximum correlations across 500 bootstrap samples for a given sample size. Black dotted lines represent the lower and upper bounds of the 95% CIs for a given sample size. The areas under the curve (AUC) for the brain-behavior associations also are displayed.

The effect sizes produced by the standard and bagging methods using the weakest edge in the functional connectome were comparable for the NIH Toolbox (2.62% < ΔAUC < 4.59%) and CBCL-externalizing (3.01% < ΔAUC < 3.35%) across the 3 low-motion racial/ethnic groups.

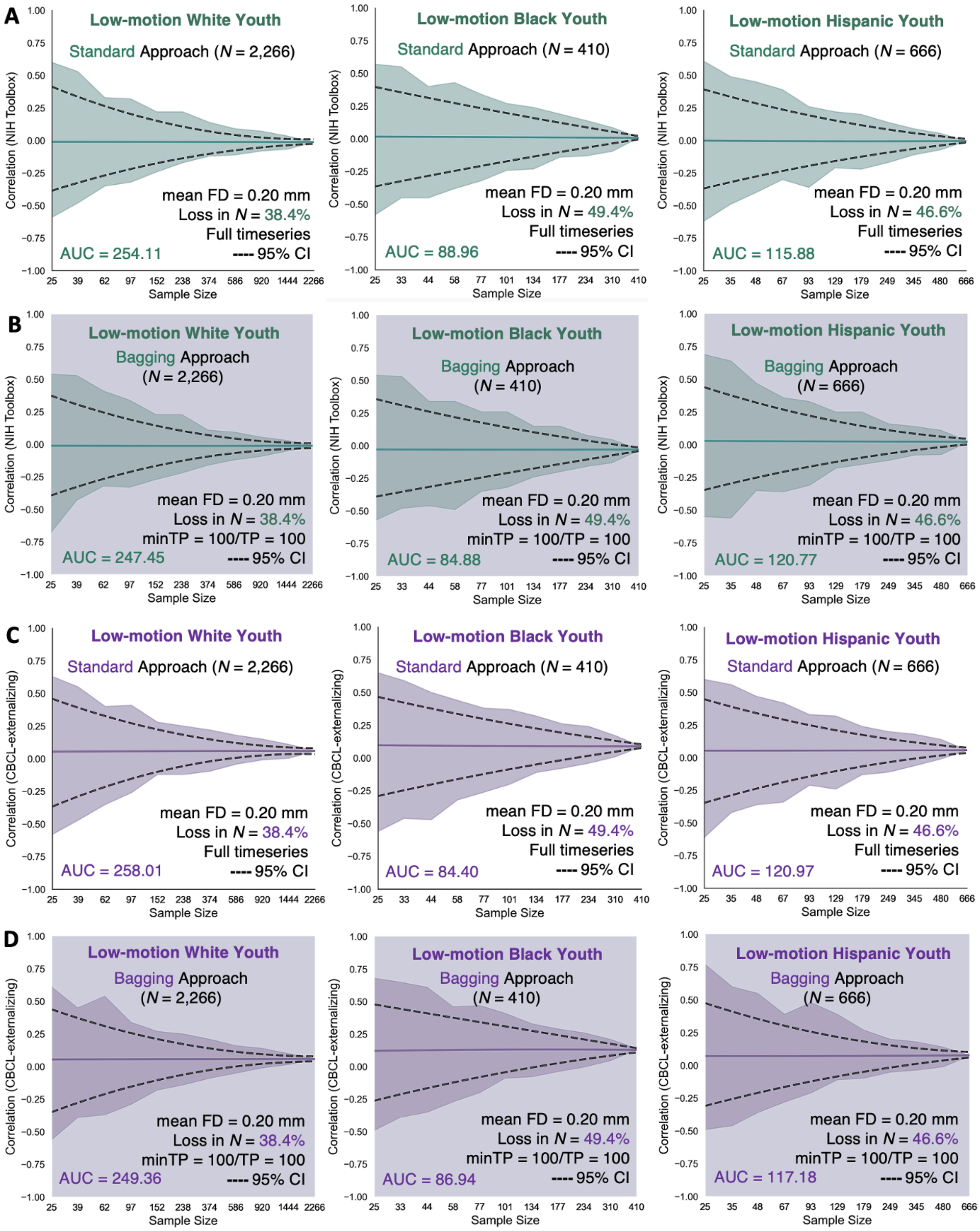


**Supplementary Figure 12. Correlations between functional connectivity and (A-B) NIH Toolbox and (C-D) CBCL-externalizing using the standard and bagging methods as a function of sample size across the 3 low-motion racial/ethnic groups obtained from the weakest edge in the functional connectome.** The respective edge that was chosen in the functional connectome shared the weakest correlation strength with the NIH Toolbox ($|R_s| = 0.042$, $P < 0.05$ FDR) and CBCL-externalizing ($|R_s| = 0.070$, $P < 0.05$ FDR). The standard method corresponded to the brain-behavior associations derived from the full fMRI timeseries of the low-motion youth with a mean FD < 0.20 mm. The bagging method corresponded to the brain-behavior associations derived from the scrubbed fMRI timeseries ranked and thresholded by their 100 least motion-corrupted timepoints ($0 < FD < 0.20$ mm) from which 100 timepoints were bootstrapped across 500 iterations to construct the functional connectivity matrices of the low-motion youth. The sample sizes were bootstrapped at 11 logarithmically-spaced $N$ intervals: White $N \in \{25, 39, 62, 97, 152, 238, 374, 586, 920, 1444, 2266\}$; Black $N \in \{25, 33, 44, 58, 77, 101, 134, 177, 234, 310, 410\}$; Hispanic $N \in \{25, 35, 48, 67, 93, 129, 179, 249, 345, 480, 666\}$. Solid teal and purple lines show the mean correlations from the 500 bootstrap samples for a given same size. Teal and purple shadings denote the minimum and maximum correlations across 500 bootstrap samples for a given sample size. Black dotted lines represent the lower and upper bounds of the 95% CIs for a given sample size. The areas under the curve (AUC) for the brain-behavior associations also are displayed.

The standard and bagged effect sizes that were obtained from the weakest edge in the functional connectome remained comparable for the NIH Toolbox (0.70% < ΔAUC < 3.42%) and CBCL-externalizing (0.05% < ΔAUC < 3.42%) across the 3 racial/ethnic groups when the high-motion youth were included.

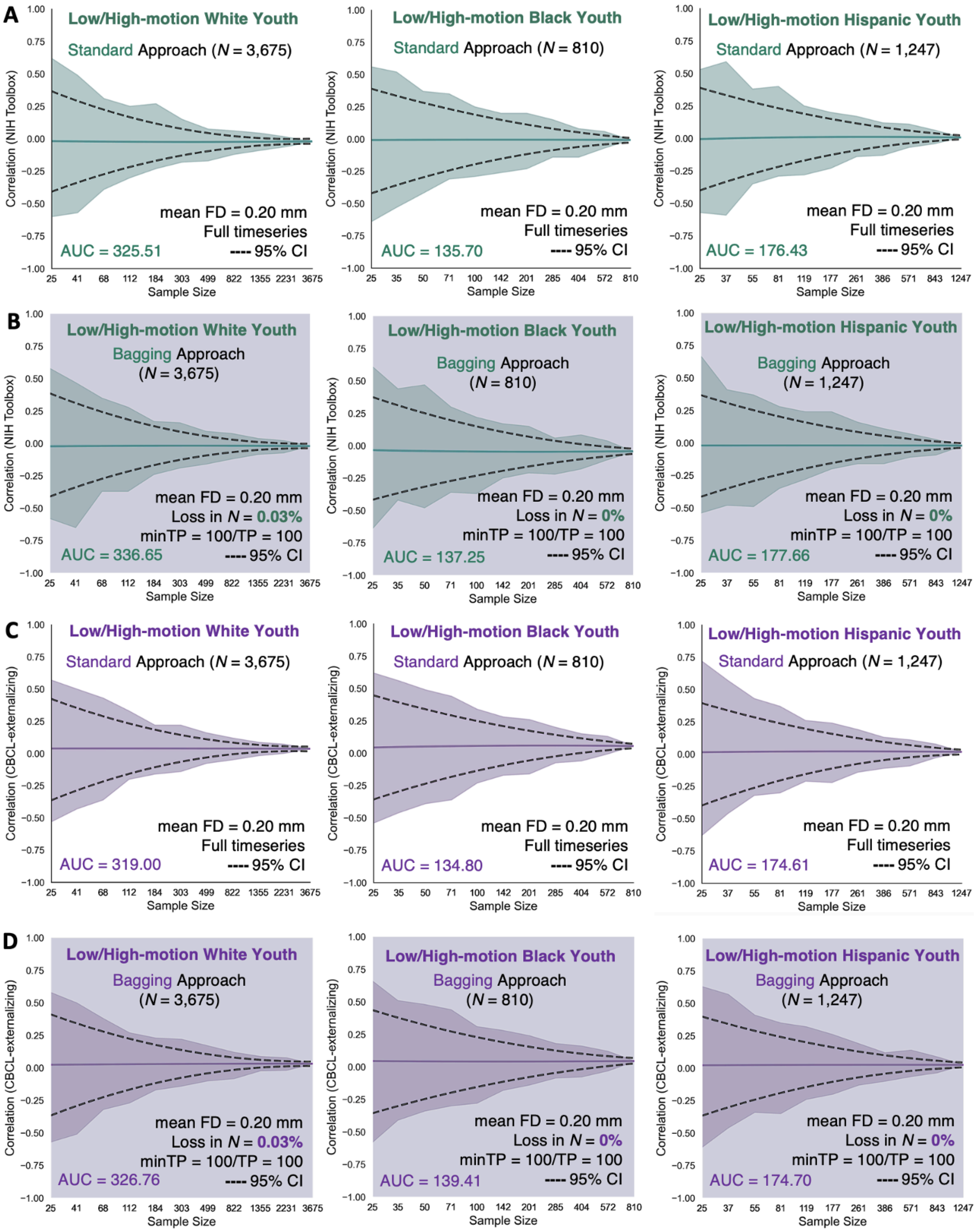


**Supplementary Figure 13. Correlations between functional connectivity and (A-B) NIH Toolbox and (C-D) CBCL-externalizing using the standard and bagging methods as a function of sample size across the 3 low/high-motion racial/ethnic groups obtained from the weakest edge in the functional connectome.** The respective edge that was chosen in the functional connectome shared the weakest correlation strength with the NIH Toolbox ($|R_s|$ = 0.042, $P$ < 0.05 FDR) and CBCL-externalizing ($|R_s|$ = 0.070, $P$ < 0.05 FDR). The standard method corresponded to the brain-behavior associations derived from the full fMRI timeseries of the low-/high-motion youth that have been retained for the analyses without imposing an initial head motion threshold of mean FD < 0.20 mm. The bagging method corresponded to the brain-behavior associations derived from the scrubbed fMRI timeseries of the low-/high-motion youth who were retained based on the assumption that they had a minimum of 100 least motion-corrupted timepoints. Their scrubbed fMRI timeseries were ranked by their lowest FD values and 100 least motion-corrupted timepoints (0 < FD < 0.20 mm) were selected from which 100 timepoints were bootstrapped across 500 iterations to construct the functional connectivity matrices of the youth. The sample sizes were bootstrapped at 11 logarithmically-spaced $N$ intervals: White $N \in \{25, 41, 68, 112, 184, 303, 499, 822, 1355, 2231, 3675\}$; Black $N \in \{25, 35, 50, 71, 100, 142, 201, 285, 404, 572, 810\}$; Hispanic $N \in \{25, 37, 55, 81, 119, 177, 261, 386, 571, 843, 1247\}$. Solid teal and purple lines show the mean correlations from the 500 bootstrap samples for a given same size. Teal and purple shadings denote the minimum and maximum correlations across 500 bootstrap samples for a given sample size. Black dotted lines represent the lower and upper bounds of the 95% CIs for a given sample size. The areas under the curve (AUC) for the brain-behavior associations also are displayed.

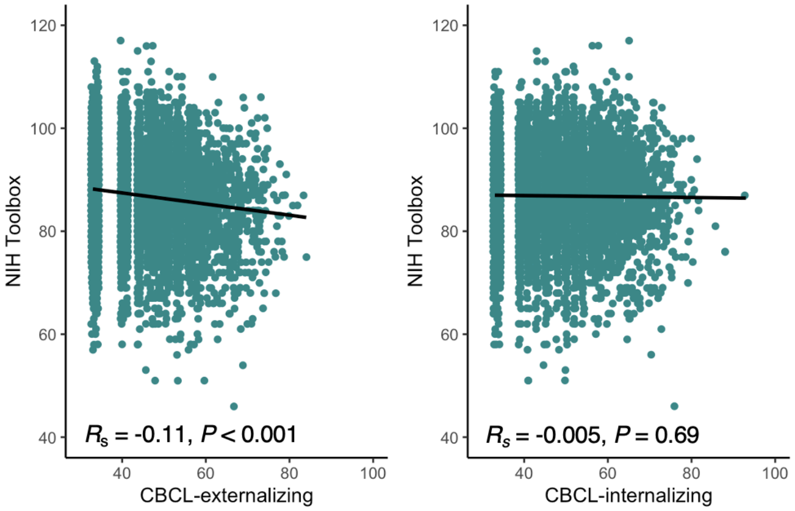


| Covariate | $R_s$ (NIH Toolbox, CBCL-Externalizing) | $R_s$ (NIH Toolbox, CBCL-Internalizing) |
|---|---|---|
| — | -0.11*** | -0.005 |
| Age (years) | -0.11*** | -0.010 |
| Site | -0.11*** | -0.002 |
| ADI[a] | -0.11*** | 0.0003 |
| Puberty[b] | -0.11*** | -0.011 |

**Supplementary Figure 14. Relationships between NIH Toolbox, CBCL-externalizing, and CBCL-internalizing while adjusting for baseline age, imaging site, area deprivation index (ADI), and pubertal status.** There was a significant relationship between NIH Toolbox and CBCL-externalizing ($R_s$ = -0.11, $P$ < 0.001) but not between NIH Toolbox and CBCL-internalizing ($R_s$ = -0.005, $P$ = 0.69). These relationships persisted after covarying baseline age, imaging site, ADI, and pubertal status. [a]The sample size was reduced from $N$ = 5,733 to $N$ = 5,457 when ADI was treated as a covariate due to missing ADI values for the youth. [b]The sample size was reduced from $N$ = 5,733 to $N$ = 5,547 when pubertal status was treated as a covariate due to missing values from the parent-report Pubertal Development Scale for the youth. The pubertal status was obtained from the male and female category subscales of the Pubertal Developmental Scale based on the youth's sex assigned at birth. ***$P$ < 0.001.

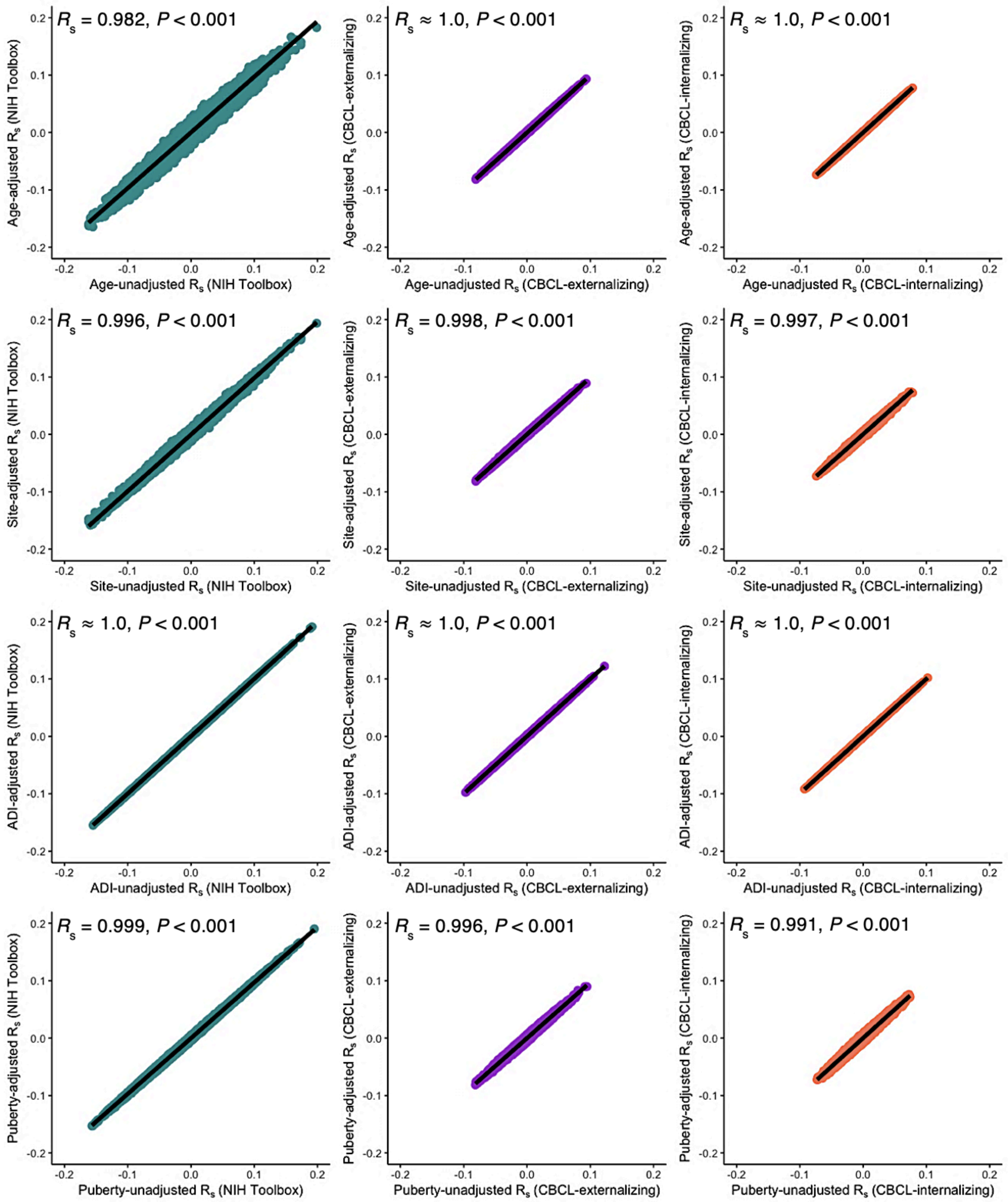


**Supplementary Figure 15. Relationships between unadjusted and adjusted edge-level correlations for baseline age, imaging site, area deprivation index (ADI), and pubertal status derived for the NIH Toolbox, CBCL-externalizing, and CBCL-internalizing from the low-motion youth.** Unadjusted edge-level correlations referred to the partial Spearman's Rank correlations between functional connectivity and the 3 behaviors while treating sex assigned at birth and mean FD as covariates across 61,776 edges in the functional connectome. Adjusted edge-level correlations referred to the partial Spearman's Rank correlations between functional connectivity and the 3 behaviors while treating sex assigned at birth and mean FD as covariates in addition to respectively adjusting for baseline age, imaging site, ADI, and pubertal status across 61,776 edges in the functional connectome. The sample size was reduced from $N$ = 3,342 to $N$ = 3,177 when ADI was covaried due to missing ADI values for the youth. The sample size was reduced from $N$ = 3,342 to $N$ = 3,247 when pubertal status was covaried due to missing values from the parent-report Pubertal Development Scale for the youth. The pubertal status was obtained from the male and female category subscales of the Pubertal Developmental Scale based on the youth's sex assigned at birth.
